# A hierarchical computational motif unifies neural dynamics across the ventral visual stream

**DOI:** 10.64898/2026.05.18.726101

**Authors:** Josh M. Wilson, Khaled Jedoui, Paolo Papale, Margaret Livingstone, Justin L. Gardner, Daniel L. K. Yamins

**Affiliations:** Department of Psychology, Stanford University; Wu Tsai Neurosciences Institute, Stanford University; Department of Computer Science, Stanford University; Department of Medicine and Surgery, Neuroscience Unit, University of Parma; Department of Vision & Cognition, Netherlands Institute for Neuroscience (KNAW); Department of Neurobiology, Harvard Medical School

## Abstract

Neural representations within individual visual cortical areas are dynamic; they evolve over tens to hundreds of milliseconds, even in response to static images. These dynamics have typically been described as area-specific phenomena with unique computational signatures. Here, we show that representational dynamics across the ventral visual stream follow a common motif: representations within each area shift over time along the same complexity axis that organizes hierarchical visual areas. The spatiotemporal signatures of this shift indicate that it is broadly distributed across the neural population, rather than concentrated in specific subpopulations. These shifts are functionally consequential, allowing for the recognition of more complex images over the course of the response. Further, we show evidence in all visual areas of a 30 ms within-area predictivity signal whose properties are consistent with local recurrence, which may be driving the representational shifts. Yet, we show that current state-of-the-art dynamic models, including one with built-in local recurrent processing, fail to recapitulate the measured neural dynamics. Together, these results reveal a common temporal motif across the ventral hierarchy, suggest local recurrence as a possible driver, and provide a concrete dynamic target for future models of the ventral visual stream.

## 1 Introduction

The primate ventral visual stream supports key perceptual behaviors through a set of hierarchical cortical areas (Ungerleider and Mishkin [1982], Goodale and Milner [1992], Felleman and Van Essen [1991], Haxby et al. [2001]). For example, population representations in these visual areas enable rapid, invariant identification of objects and categories, a computational objective central to primate vision and behavior (Hung et al. [2005], DiCarlo and Cox [2007], DiCarlo et al. [2012], Logothetis and Sheinberg [1996], Tanaka [1996]). More successfully than perhaps any other cortical system, the ventral visual stream has served as a testbed for theories of how the brain efficiently and effectively encodes sensory input into cortical population representations (Hubel and Wiesel [1959, 1962], Carandini and Heeger [2012], Olshausen and Field [1996]).

At the level of temporally averaged responses, the ventral visual hierarchy is well-described by purely feedforward models that unify its areas along a single axis of representational complexity that increases with hierarchical position. V1 neurons are selective for oriented edges and local spatial frequencies (Hubel and Wiesel [1968], Simoncelli and Olshausen [2001], Carandini et al. [2005]); intermediate areas encode nonlinear combinations of these features, including texture statistics, boundary curvature, and shapes (Freeman et al. [2013], Pasupathy and Connor [1999, 2001], Gallant et al. [1993]); inferotemporal cortex (IT) encodes further nonlinear combinations of mid-level features in the form of complex object properties (Tanaka [1996], Gross et al. [1972], Desimone et al. [1984], Jagadeesh and Gardner [2022]). Across these areas, selectivity and tolerance to identity-preserving transformations both increase with hierarchical position (Ito et al. [1995], Logothetis and Sheinberg [1996], Rust and DiCarlo [2010]). Purely feedforward models recapitulate this progression: alternating template-matching and pooling operations reproduce the increasing complexity and invariance of the ventral pathway (Riesenhuber and Poggio [1999]), and deep networks optimized for object recognition or other objectives develop layered representations that predict neural responses at corresponding hierarchical stages (Yamins et al. [2014], Khaligh-Razavi and Kriegeskorte [2014], Yamins and DiCarlo [2016], Cadena et al. [2019], Gü çlü and van Gerven [2015], Zhuang et al. [2021]). This success, established on neural representations averaged over the full course of the response, has reinforced a view in which representational complexity is fixed by hierarchical position – and, implicitly, that this fixed correspondence is the right description of the response throughout its time course.

However, representations within visual areas are dynamic: even in response to static stimuli, representations evolve over the course of the response in ways that purely feedforward models, by construction, cannot reproduce (Kietzmann et al. [2019], Lamme and Roelfsema [2000], Kreiman and Serre [2020]). Time-resolved analyses of macaque V4 and IT reveal information transfer that varies temporally and semantically, with recurrent decoders extending across time outperforming temporally local decoders (Anthes et al. [2026]). Images that feedforward networks have trouble discriminating require additional processing time in IT to reach behaviorally sufficient population codes (Kar et al. [2019]). Task-optimized recurrent networks predict V4 and IT response trajectories that their feedforward counterparts cannot, across the full stimulus presentation (Nayebi et al. [2018, 2022]). How these dynamics actually unfold within individual cortical areas has been the subject of decades of investigation.

These dynamics have typically been documented separately across visual areas and characterized as distinct, area-specific phenomena. For example, V1 neurons have been shown to sharpen their tuning for orientations and spatial frequencies over the course of the response (Ringach et al. [1997], Mazer et al. [2002], Bredfeldt and Ringach [2002]). V4 responses convey information about multipart configurations and curvature later than they do simpler features (Brincat and Connor [2006], Yau et al. [2013]). IT neurons initially encode global category information and shift toward fine-grained identity as the response unfolds (Sugase et al. [1999], Matsumoto et al. [2005], Shi et al. [2026]). Each phenomenon was discovered separately, in a different area, with a different stimulus paradigm, contributing to an area-specific interpretation of seemingly different dynamics.

Here, we use a model-driven technique to precisely quantify temporal deviations from feedforward computations, identifying a common dynamical form across the ventral visual stream: within each area, representations shift over time along the same axis of representational complexity by which hierarchical visual areas are organized. Using simultaneous large-scale multi-unit macaque recordings spanning V1, V4, and IT (Papale et al. [2025]) together with a feedforward model (DINOv2, Oquab et al. [2023]) whose layered representations match the ventral hierarchy at a level approaching cross-animal consistency, we tracked how each area’s correspondence to the model evolved over the course of the response. The network layer that best predicted each area progressively deepened over time, in violation of the feedforward prediction of a fixed mapping between network depth and cortical area. These shifts were common to V1, V4, and IT.

We further show that this shared form reflects a shared underlying computational process operating within each area. The shift extended to the level of individual electrodes; it was broadly distributed across each area rather than concentrated in subpopulations. Moreover, the shift was not explained by staggered recruitment of late-responding electrodes. These shifts had functional consequences, allowing for the decoding of more complex images over time. We go on to show that the shifts were accompanied by a 30 ms within-area predictivity enhancement: same-area activity 30 ms earlier predicted current responses better than activity at shorter offsets, with a short-range spatial profile and a time course that tracked the complexity shift itself. While local recurrent processing within each area provides a parsimonious account of these observations, we show that state-of-the-art dynamic neural network models – including one with explicit local recurrent circuitry – do not capture these shifts, providing a target for future modeling efforts.

## 2 Results

To study the temporal dynamics of representations in the ventral visual stream, we used macaque multi-unit activity (MUA) data from the THINGS Ventral Stream Spiking Dataset (TVSD; Papale et al. [2025]). The dataset provides millisecond-resolution recordings from V1, V4, and IT in two macaques viewing 22,348 static images from the THINGS dataset (Hebart et al. [2019]; see original publication and Methods for full details).

We leveraged the temporal resolution and multi-area coverage of TVSD recordings to characterize how representational complexity evolves within individual cortical areas and to identify the computations underlying these dynamics. We first establish that a feedforward model (DINOv2, Oquab et al. [2023]) parallels the ventral stream at a level approaching cross-animal consistency when evaluated on representations averaged over the course of the neural response (Fig. 1). These results validated the network hierarchy as a complexity axis akin to the ventral visual hierarchy. We then track how each area’s correspondence to this network hierarchy changed at high temporal resolution (10 ms bins) to characterize how representations within each area evolved over the course of the response (Fig. 2). To characterize how these shifts are composed across the population, we examine the distribution of representational complexity shifts across the recorded population (Figs. 3–4). To determine the functional significance of the complexity shifts, we show that images requiring deeper network representations were decoded later from neural activity as representations became more complex (Fig. 5). We then examine the temporal signature of these shifts and find a within-area predictivity enhancement consistent with local recurrent processing (Fig. 6). With this candidate mechanism in mind, we show that state-of-the-art dynamic neural network models – even one with explicit local recurrent circuitry – did not capture the same pattern of shifts as the brain (Fig. 7).

**Figure 1:**
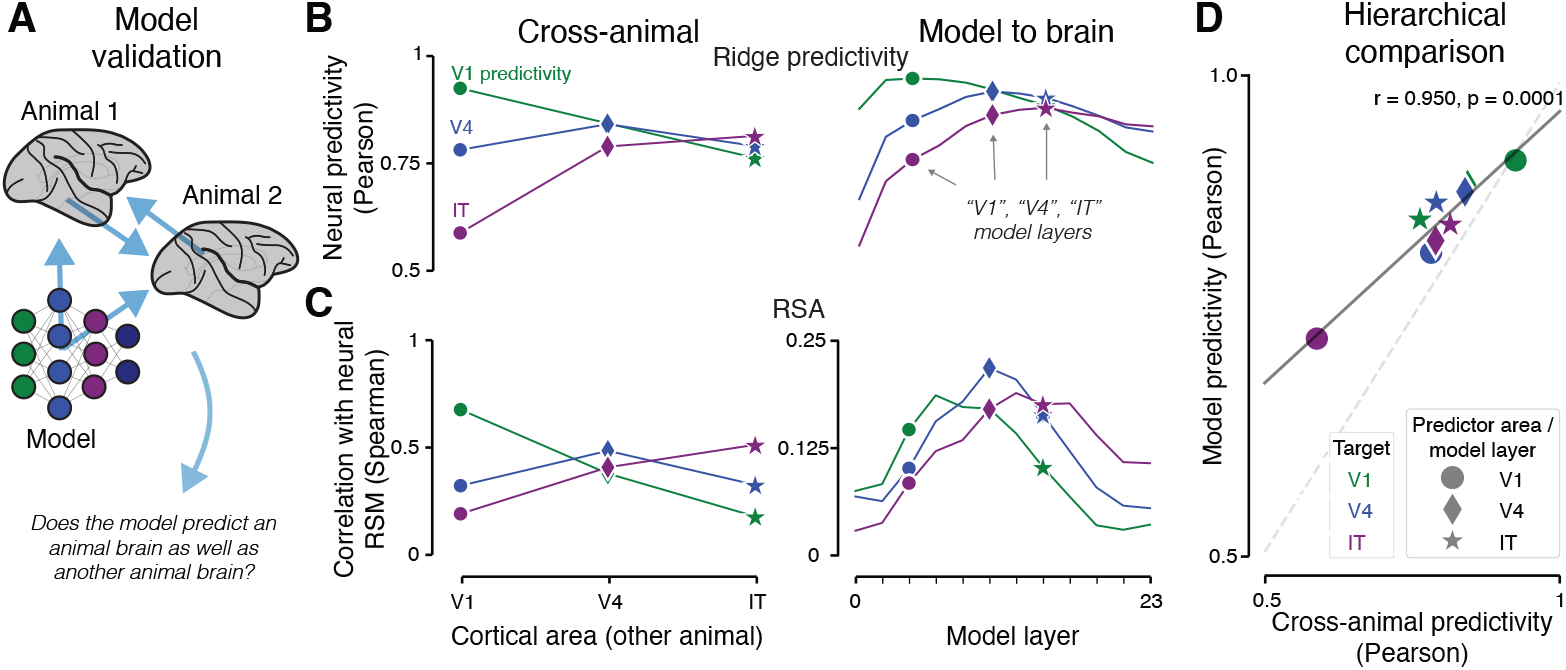
Temporally averaged cross-animal and model-animal mapping patterns are highly aligned along the ventral visual stream. (A) Validation framework. (B) Ridge regression predictivity for cross-animal (left) and model-to-brain (right) predictions. Each point shows the median predictivity (Pearson correlation on the held-out test set) across both monkeys for all electrodes in a target area (colors). Shapes mark the most-predictive layer for each target area. (C) Representational similarity analysis using both cross-animal and model-to-brain approaches. Conventions as in (B): points represent the mean Spearman correlation between predictor and target representational similarity matrices across the two monkeys. (D) Model predictivity versus cross-animal predictivity for each of nine predictor-to-target area mappings. For each pair, the model layer that best predicts the predictor area is used to predict the target area. Color indicates target area; shape indicates predictor area and corresponding model layer (same areas/layers as (B)). Solid line shows linear fit of model predictivity on cross-animal predictivity.

**Figure 2:**
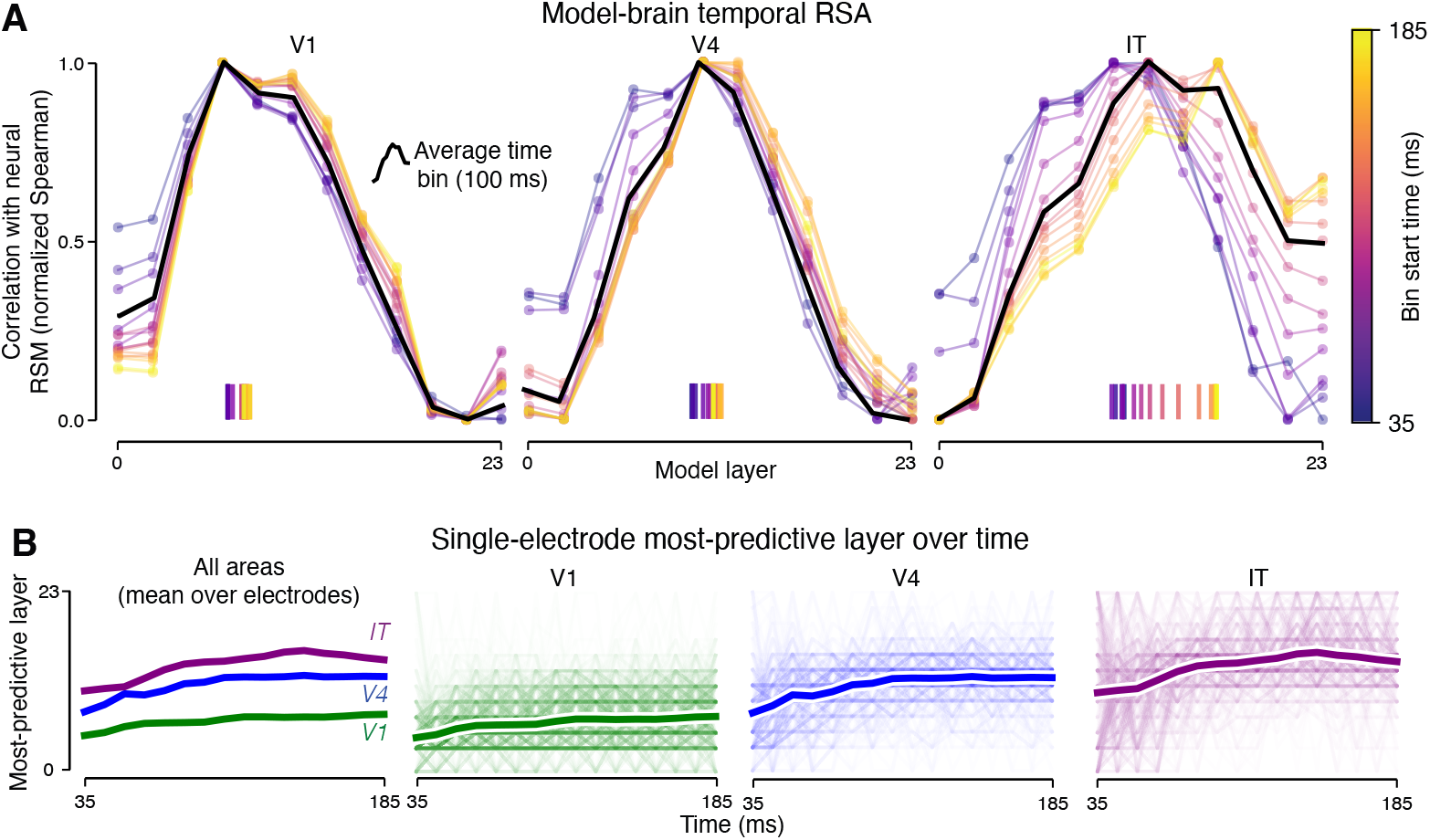
Fine-grained temporal dynamics of model-brain alignment shift toward deeper network layers over the course of the response, at both the population and individual-electrode level. (A) Model-brain temporal RSA. For each 10 ms time bin (color) and cortical area, the Spearman correlation between the neural population RSM and each model layer RSM is shown, normalized to span 0-1 within each time bin. The black line shows the same correlation profile computed from the time-averaged (100 ms window) response, for reference. Tick marks along the bottom indicate the softmax-weighted center of mass of each time bin’s curve across layers. (B) Most-predictive model layer over time for all electrodes in V1, V4, and IT, calculated as per-electrode argmaxes from ridge regression. Thick lines show the mean best layer for each time bin across electrodes within each area; colors denote visual area. Right panels: thin lines show individual electrodes; thick line repeats the area mean.

**Figure 3:**
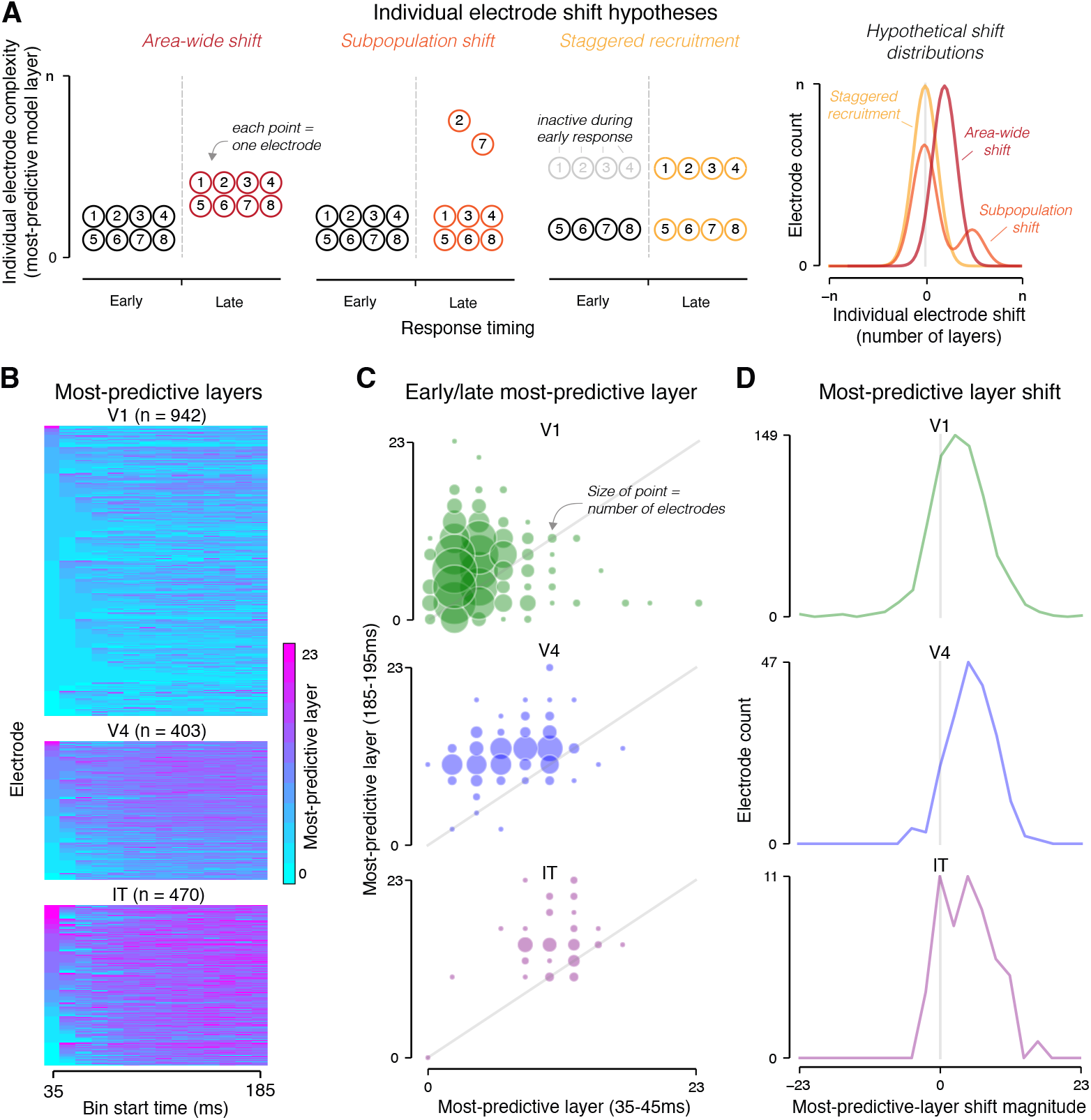
Complexity shifts are distributed broadly across electrodes rather than concentrated in a subpopulation. (A) Left, schematic of three hypotheses for how a population-level shift could be composed at the single-electrode level. Each point represents one electrode, plotted by response timing (early vs. late) and complexity (most-predictive model layer). Right, distributions of individual shift magnitudes predicted under different hypotheses. (B) Most-predictive layers of individual electrodes over different time bins. Count indicates number of electrodes with an average split-half reliability *>*0.2 across time bins. Color denotes most-predictive layer. (C) Scatter plot of the most-predictive layer at 35-45 ms and 185-195 ms for each electrode in each visual area. Point size indicates number of electrodes. In addition to the aforementioned reliability cutoff, only electrodes with reliable signal in the earliest time bin are plotted (*>*0.2 split-half reliability). (D) Amount of shift in individual electrodes from first to last time bin, calculated as the number of layers above (positive) or below (negative) the diagonal in (C).

**Figure 4:**
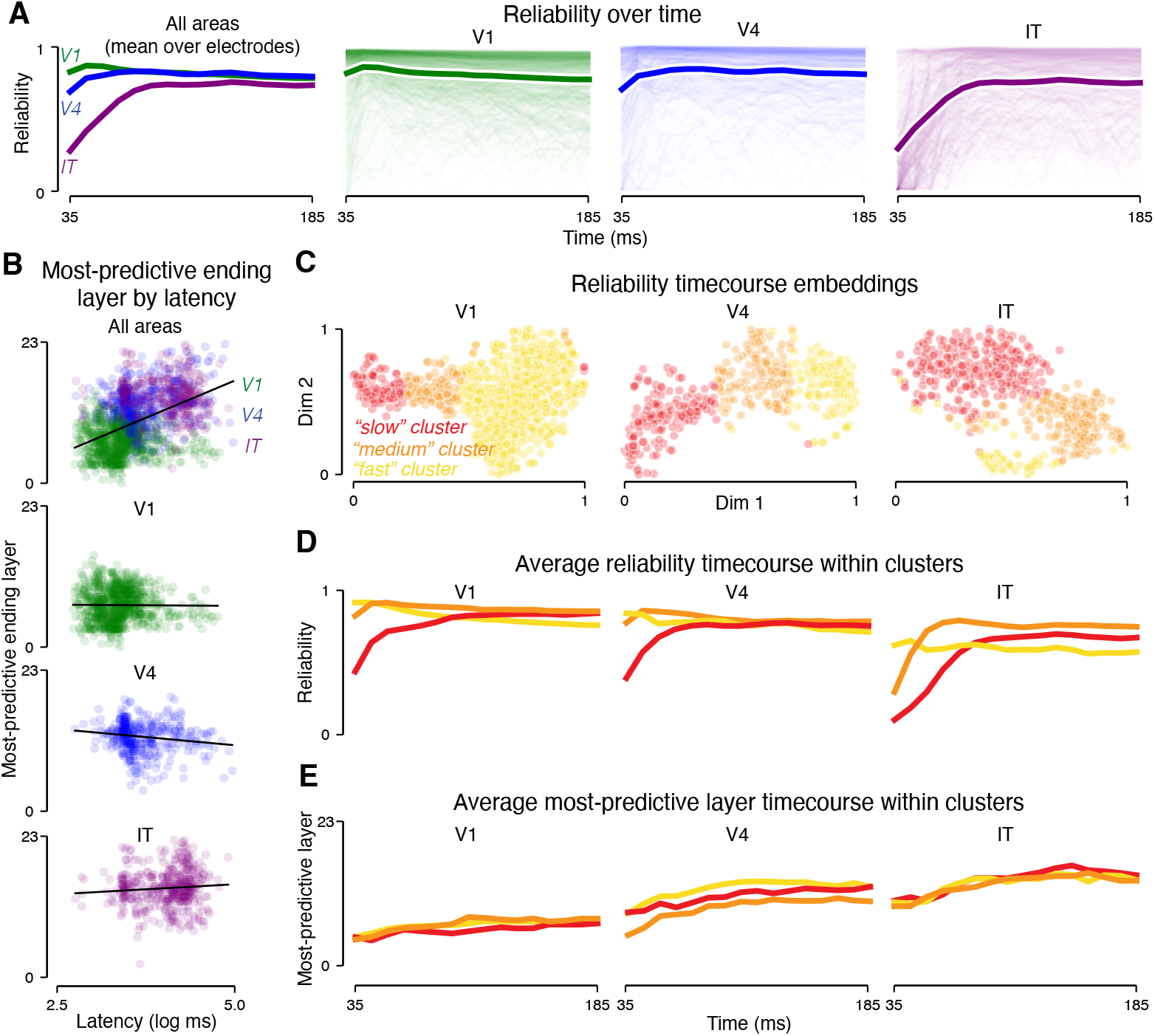
Despite heterogeneity in response latencies, electrodes follow similar complexity trajectories within each area, and latency and complexity are dissociable. (A) Split-half reliability over time for all electrodes in V1, V4, and IT. Thick lines show the mean reliability across electrodes in an area for each time bin; thin lines show individual electrodes. Colors denote visual areas. (B) Individual electrodes plotted by response latency and most-predictive ending layer. Latency estimates were taken from Papale et al. [2025] and averaged over all recording days. Most-predictive ending layer was calculated as the mean most-predictive layer across the last five time bins, rather than just the last time bin. Top row: all electrodes pooled across areas. Bottom rows: individual areas. Lines show best linear fit. (C) Two-dimensional t-SNE embeddings of reliability over time, embedded separately for the three visual areas. Points are individual electrodes. Colors denote three k-means clusters, found in the native space and overlaid on the embeddings (red: slow, orange: medium, yellow: fast). (D) Mean split-half reliability over time of electrodes in each cluster. (E) Mean most-predictive layer over time for electrodes in each cluster.

**Figure 5:**
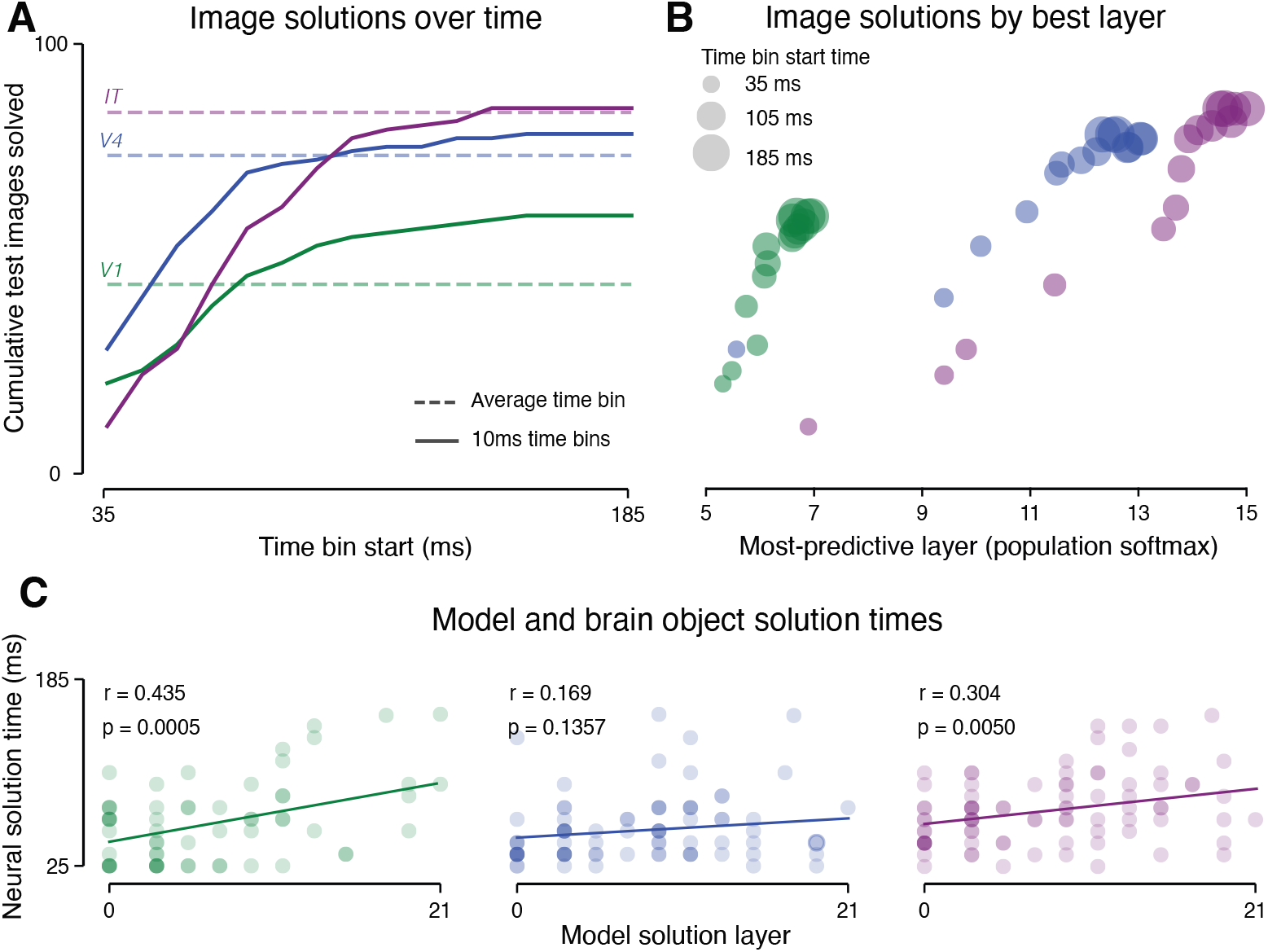
Image-level classification accuracy patterns correlate with complexity shifts, and images requiring deeper network features are decoded later. (A) Cumulative number of test images solved over time by logistic regression classifiers trained on neural data from each area (color: green, V1; blue, V4; purple, IT). Solid lines show classifiers trained on 10 ms time bins; dashed lines show classifiers trained on the time-averaged (100 ms window) response. An image is considered solved when its correct category falls within the top 5 predictions for two consecutive time bins. (B) Cumulative images solved versus the most-predictive model layer (softmax-weighted center of mass from ridge regression) for each area at each time bin. Bubble size indicates time bin (larger = later); color indicates area. (C) Neural object solution time versus network solution layer for individual images. Each dot is one test image (darker dots indicate overlapping images); color indicates area. Neural solution time is the time bin at which the image is first solved by the neural classifier. Network solution layer is the model layer at which a logistic regression classifier first correctly classifies the image. Lines show linear fits.

**Figure 6:**
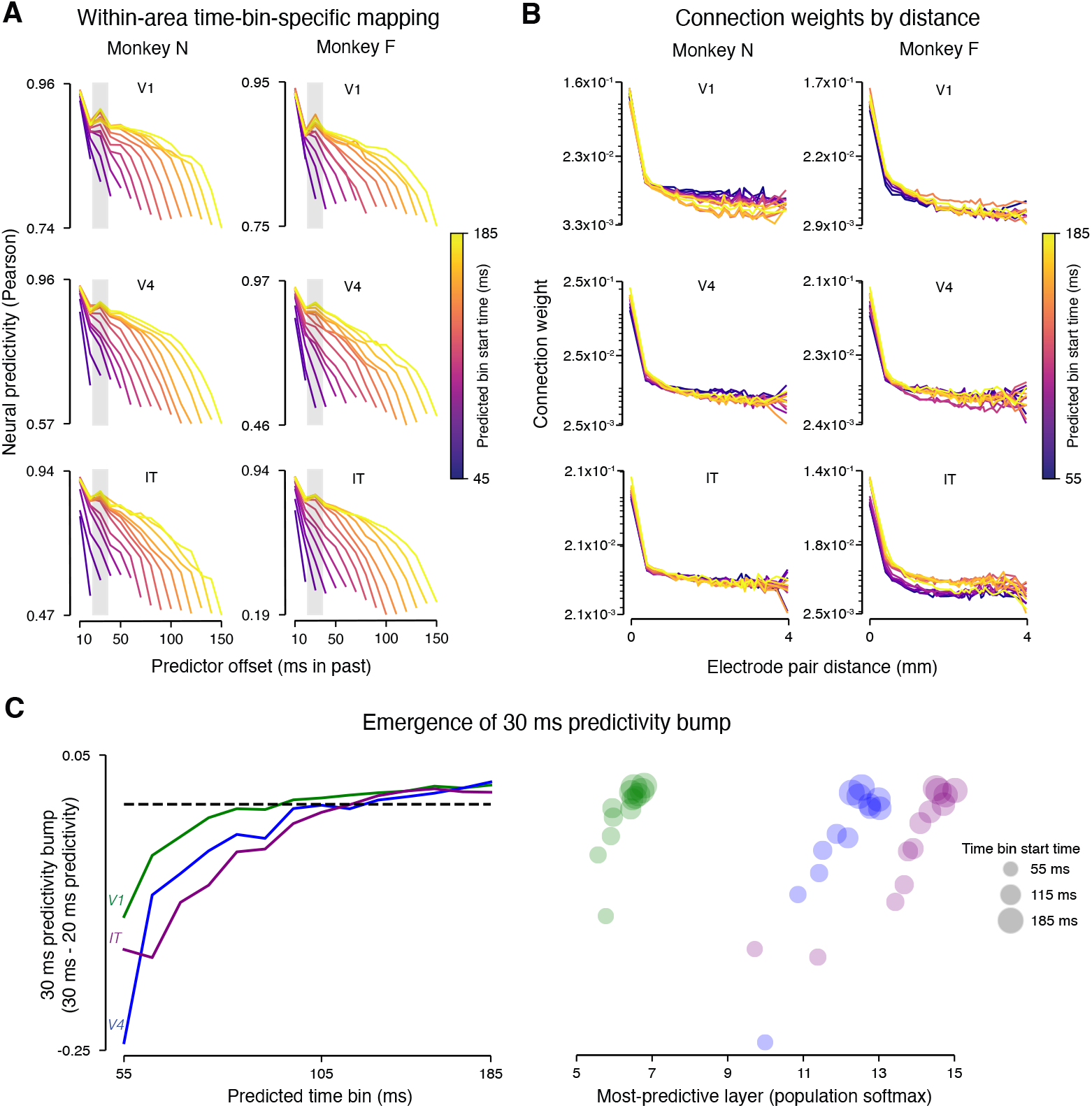
Time-lagged predictivity patterns reveal a non-monotonic enhancement at 30 ms that is spatially local and co-emerges with the complexity shifts. (A) Within-area ridge regression predictivity between responses in different time bins. For each predicted time bin, predictivity is shown as a function of the temporal offset between predictor and target bins. Rows show V1, V4, and IT; columns show Monkey N and Monkey F. Shaded columns show 30 ms offset. (B) Ridge regression coefficient magnitude as a function of inter-electrode distance for the 30 ms offset mapping in (A). Distances are computed within arrays; values are averaged across arrays. (C) Emergence of the 30 ms predictivity bump. Left: magnitude of the predictivity bump at 30 ms (relative to adjacent 20 ms offset) for each predicted time bin and area. Right: the 30 ms bump magnitude plotted against the most-predictive model layer at each time bin for each area. Bubble size indicates predicted time bin (larger = later).

**Figure 7:**
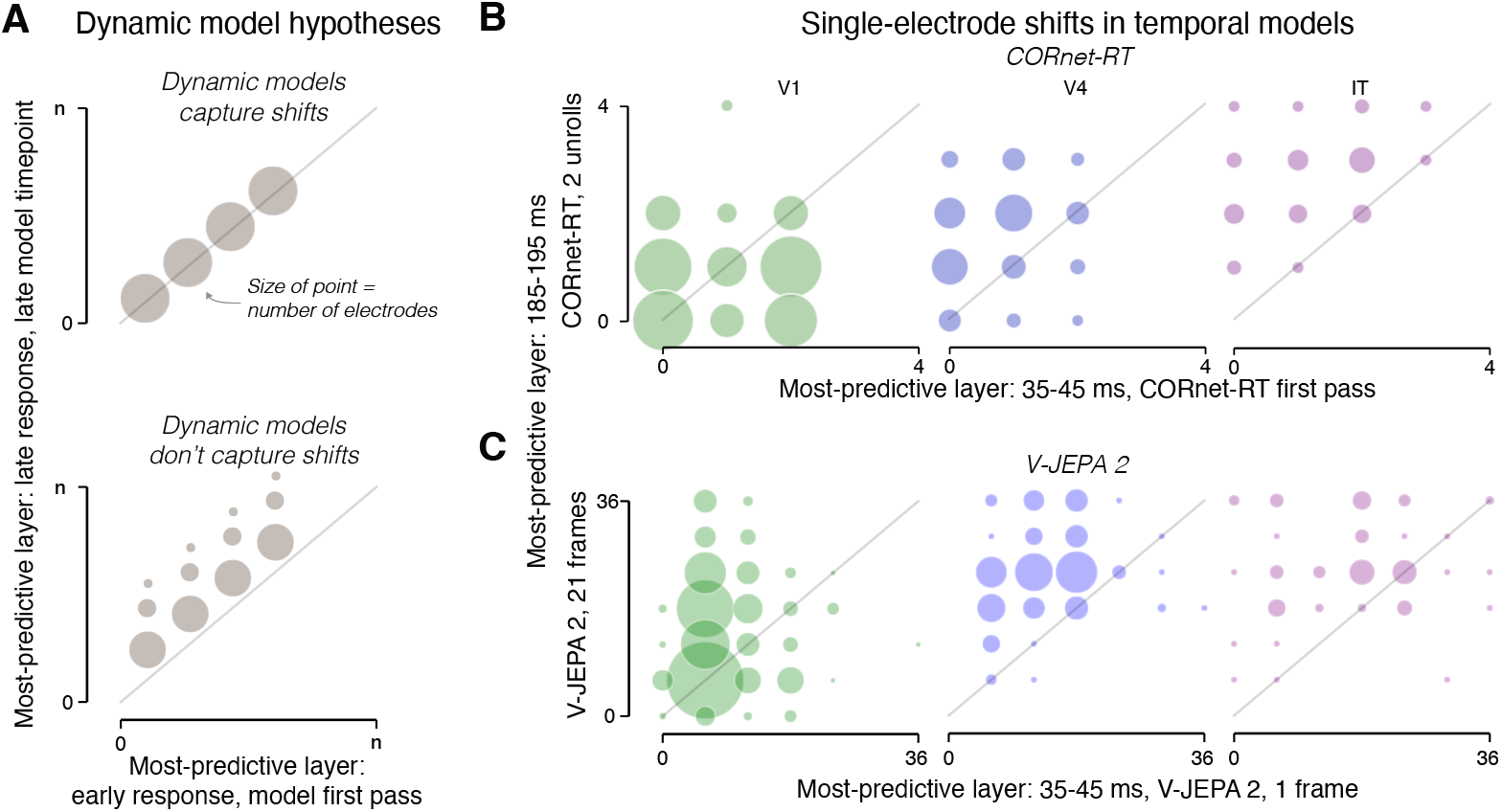
Two classes of dynamic neural network models do not capture representational shifts. (A) Schematic of expected outcomes for the per-electrode shift analysis. Conventions similar to Fig. 3C. (B) Most-predictive CORnet-RT layer (first pass) at 35–45 ms versus most-predictive CORnet-RT layer (after two additional biological unrolls) at 185–195 ms for each electrode. Point size indicates number of electrodes. (C) Most-predictive V-JEPA 2 layer at 35–45 ms (1-frame features) versus 185–195 ms (21-frame features, retaining only the tokens corresponding to the most recently tokenized input) for each electrode. Conventions as in (B).

### 2.1 A feedforward model captures temporally averaged neural response patterns at high fidelity across the ventral hierarchy

Given a model of the brain whose hierarchy parallels that of the ventral stream, representational dynamics within each cortical area can be quantified as the change over time in how the area’s representations correspond to the fixed model’s hierarchy. We start by establishing such a model. Cross-animal predictivity provides the empirical standard for evaluating whether a candidate model’s hierarchy parallels the ventral stream’s (Thobani et al. [2025]) (Fig. 1A). A sufficient model reproduces not just the predictivity of each area, but the full pattern of graded inter-area relationships at magnitudes matching cross-animal predictions. A model that passes this test provides the calibrated hierarchy needed to track representational dynamics in the time-resolved domain. Here, we show that a feedforward model (DINOv2) meets this standard on time-averaged responses (100 ms windows) across V1, V4, and IT.

Cross-animal mappings revealed clear hierarchical organization when averaging over 100 ms response windows. We summarized each electrode’s response as the average MUA over a 100 ms window (starting at 25, 50, and 75 ms post-stimulus for V1, V4, and IT, respectively), then used ridge regression to predict image-evoked responses from responses in different areas of the other animal (Fig. 1B, left). Each area was best predicted by its counterpart in the other animal (V1: r = 0.925, V4: r = 0.842, IT: r = 0.814, Pearson correlation between predicted and actual test-set responses) (Fig. 1B, left), with predictivity declining with hierarchical distance. Representational similarity analysis (RSA, Kriegeskorte et al. [2008]) confirmed the same hierarchical ordering without a fit linear mapping (Fig. 1C, left).

Model-to-brain mappings exhibited the same predictivity patterns as brain-to-brain mappings at comparable magnitudes. Predicting responses in each target area from features extracted at successive layers of the network yielded a depth-dependent profile in which earlier layers best predicted V1, intermediate layers V4, and later layers IT (Fig. 1B, right). Peak model-to-brain predictivity approached or exceeded cross-animal predictivity in all three areas (V1: r = 0.917, V4: r = 0.887, IT: r = 0.849). RSA again confirmed the same depth-dependent alignment without a fit mapping (Fig. 1C, right).

Moreover, the model reproduced the full pattern of inter-area predictivity, not just same-area predictions. To test whether the model could stand in for another brain across the full set of area comparisons, we selected the model layer that best predicted each predictor area and used that layer’s features to predict each target area, mirroring the nine cross-animal mappings. Model predictivities for these nine mappings were tightly correlated with the corresponding cross-animal predictivities (Fig. 1D; r = 0.950, p = 0.0001).

The model thus reproduced not only the hierarchical ordering but the graded pattern of inter-area relationships at magnitudes expected of another brain. This agreement between cross-animal and model-brain mappings, both in predictive magnitude and the full pattern of inter-area relationships, establishes the model as a hierarchically aligned model of the time-averaged ventral stream representations, and defines a complexity axis along which individual electrodes can be localized within the static hierarchy: successive layers of the model.

### 2.2 Fine-grained temporal dynamics systematically diverge from feedforward predictions, becoming more complex over time

Having established a temporal-average baseline with a brain-like feedforward model, we employed the model to measure systematic deviations from the feedforward structure at finer-grained temporal resolution. Feedforward models of the ventral stream predict that the correspondence between network depth and cortical area should be fixed: early layers map best to V1, intermediate layers to V4, and late layers to IT, regardless of when in the response the mapping is evaluated. We tested this prediction by applying time-resolved versions of both model-brain and cross-animal RSA to track how these mappings changed over the course of the evoked response. To do so, we computed population representational similarity matrices (RSMs) from MUA responses averaged over 10 ms bins. For model-brain comparisons, we compared each time bin’s RSM to RSMs from successive model layers, normalizing each bin’s correlation profile to span 0 to 1 to isolate changes in the depth profile from changes in overall magnitude driven by time-varying reliability. Unnormalized profiles show the same shift (Supplementary Fig. 4). For cross-animal comparisons, we compared each time bin’s RSM in one animal to the fixed time-averaged RSM of each area in the other animal, rather than comparing time-resolved RSMs simultaneously in both monkeys, applying the same per-time-bin normalization.

Representations within each cortical area progressively shifted toward greater complexity over the course of the response, as evidenced by time-varying alignment with both model layers and other cortical areas. Time-resolved model-brain RSA revealed that in all three areas, the center of the normalized correlation curve shifted systematically toward later model layers as time progressed, deviating from the time-averaged baseline (Fig. 2A). The same pattern was evident in other feedforward models besides DINOv2 (Supplementary Fig. 3). Cross-animal comparisons confirmed the same temporal progression without reference to an artificial network: in all three areas, representational geometry shifted over time toward higher areas in the other monkey (Supplementary Fig. 1). Time-resolved ridge regression encoding models of individual electrodes yielded a similar pattern (Supplementary Fig. 2A).

Having established these shifts at the population level, we show that they held at the individual electrode level. To do so, we identified – for each electrode and each time bin – the model layer that best predicted that electrode’s response (via ridge regression), and tracked how this best-predicting layer changed over time. In all three areas, individual electrodes shifted toward later model layers over time, following area-characteristic trajectories (Fig. 2B). Individual electrode trajectories showed variance around the area means but followed the same upward trend (Fig. 2B, thin lines; Supplementary Fig. 5A). V1 electrodes were best predicted by early layers but shifted toward intermediate layers (mean best layer across individual electrodes = 4.39 at 35 ms, 7.05 at 185 ms); V4 and IT electrodes started at intermediate layers and shifted further toward later layers (V4: 7.44 to 11.98; IT: 10.14 to 14.15, mean best layers, 35 ms to 185 ms).

Together, both measures revealed a simple, systematic deviation of neural responses from a purely feedforward prediction in all three visual areas: representations within each area shifted over time toward greater complexity along the same axis that distinguishes hierarchical stages in the ventral stream.

### 2.3 Individual-electrode dynamics implicate a population-wide process rather than specialized subpopulations

Using the per-electrode predictions, we characterized how the population-level shifts were composed at the single-electrode level. Area-level complexity shifts can arise from very different distributions of single-electrode contributions (Fig. 3A, left). For example, many electrodes may shift modestly in concert (an “area-wide shift”), or a small subset of electrodes may shift dramatically while most stay static (a “subpopulation shift”). Alternatively, the population complexity could shift without a shift in any individual electrode due to “staggered recruitment” of more complex electrodes over the course of the response. To distinguish between these possibilities, we measured the distribution of the shifts in the most-predictive model layer (maximum ridge predictivity, Pearson correlation) across individual electrodes between early and late time bins. Each shift hypothesis makes a concrete prediction for the distribution of individual-electrode shifts over time (Fig. 3A, right).

Population-level shifts were driven by shifts in many individual electrodes rather than large shifts in a few. These dynamics were visible from the most-predictive-layer timecourses (Fig. 3B). To quantify the amount of shift for each electrode, we compared each electrode’s most-predictive layer early in the response (35–45 ms) to its most-predictive layer late in the response (185–195 ms). This analysis revealed that the majority of electrodes in all three areas shifted toward later layers over time (Fig. 3C). The distribution of per-electrode shift magnitudes was centered on a modest positive value in all three areas (a shift of a few layers) with similar spread (Fig. 3D). Per-electrode shifts were significantly greater than zero in every area (one-sided t-test, p *<* 0.001). This pattern indicates that the representational complexity shift is broadly distributed across the recorded population, consistent with an area-wide mechanism.

### 2.4 Complexity shifts are not due to staggered temporal recruitment

Another possible mechanism driving the observed population-level complexity shifts is the staggered recruitment of more complex electrodes. If late-responding electrodes encode more complex visual features, their gradual recruitment could produce a population-level complexity shift without any individual electrode changing its tuning. The per-electrode analyses in Fig. 3C and 3D required reliable responses in both the earliest and latest time bins to ensure that layer assignments reflected meaningful signal rather than noise, and therefore excluded the late-responding electrodes that a staggered-recruitment account would invoke. Moreover, substantial response timing heterogeneity within each visual area (Fig. 4A) suggests that staggered recruitment may influence population-level complexity shifts. To determine if staggered recruitment was a significant driver of population complexity shifts, we compared representational complexity trajectories across electrodes with distinct timing profiles within each area (Fig. 4).

While we found a strong relationship between response latency and electrode complexity across areas, we found little evidence of a relationship within individual areas. When electrodes were pooled across areas, electrodes with later response latencies were better-predicted by later layers of the model (Fig. 4B, top; slope = 4.57, p *<* 0.001), but within each individual area this relationship was near zero or even slightly negative (Fig. 4B, bottom; slopes = -0.07, -0.99, 0.61; p = 0.801, 0.002, 0.090 for V1, V4, IT). This dissociation – a Simpson’s paradox driven by hierarchical differences across areas rather than within-area variation – suggests that staggered recruitment of electrodes does not drive the measured population-level representational complexity shifts.

Further, despite the substantial differences in reliability over time, electrodes within each area followed similar representational complexity trajectories. We clustered electrodes within each area into three groups based on their z-scored split-half reliability timecourses using k-means (Fig. 4C), capturing slow, medium, and fast timing profiles (red, orange, and yellow clusters). The resulting clusters reflected differences in temporal dynamics: within IT, for example, one cluster showed rapid onset and sustained reliability while another ramped slowly over the full response period (Fig. 4D). Despite these pronounced timing differences, the three clusters within each area showed highly similar best-layer trajectories over time (Fig. 4E) – early-responding and late-responding electrodes alike underwent comparable shifts toward later network layers.

### 2.5 Complexity shifts track image-level neural decoding ability

One functional hypothesis for why representations shift over time is that the shift enables the recognition of images that are not classifiable from the purely feedforward pass (Kar et al. [2019]). This hypothesis makes two concrete predictions in our data. First, classification performance should track representational complexity over time – more images should be solved as the representations shift. Second, images requiring more complex representations should be classified later over the course of the response. To test these hypotheses, we trained logistic regression classifiers to decode image category from the time-resolved neural population activity in each area, pooling electrodes across both monkeys, and also from features in sequential layers of a neural network model (DINOv2) (Fig. 5).

All three visual areas solved more images over time, tracking the concurrent representational complexity shifts and plateauing as representations stabilized. All three areas continued to solve additional images over the course of the response, with V4 and IT ultimately solving more images than V1 (Fig. 5A) (60, 79, and 85 out of 100 images cumulatively solved by V1, V4, and IT). Within each area, later time bins were associated with both more solved images and deeper most-predictive model layers, and the rate of newly solved images leveled off as representations approached their final complexity level (Fig. 5B). Importantly, the shift in representational complexity over time better explains the increase in solved images than increases in signal split-half reliability, which peaked early and – in V1 and V4 – declined as visual cortex was still solving new images (Supplementary Fig. 6).

Images that required deeper network representations to classify were solved later from neural activity, suggesting that the representational complexity shift reflects a progressive construction of the complexity that specific stimuli demand. For each image, we independently determined its neural object solution time (OST) and the network layer in which it was first classified correctly. Across all three areas, images requiring deeper network layers tended to be solved later in the neural response (Fig. 5C) (r = 0.435, 0.169, 0.304 in V1, V4, and IT), though this relationship was significant only in V1 and IT (p *<* 0.01 for V1 and IT; p = 0.136 in V4). While statistically weak, the V4 effect was in the expected direction; the weaker statistical reliability likely reflects both V4’s smaller recorded population (7 recorded arrays, versus 9 and 16 in IT and V1 across both monkeys) in conjunction with the classifier construction, which trains on only 12 examples per category across the 100 test-set classes. The sensitivity of the OST metric is also reflected in the low cross-animal correlation of OSTs (Supplementary Fig. 7).

Our results suggest that the complexity shifts within each area are not epiphenomenal – that representations are progressively constructed to meet the demands of specific stimuli. Images that required deeper network representations to classify were decoded later from neural activity, linking the temporal dynamics to stimulus-specific processing demands.

### 2.6 A 30 ms within-area predictivity signal exhibits the spatiotemporal properties of local recurrence

Having established the breadth, dissociability, and functional relevance of the shifts, we next looked for evidence of local recurrent processing, which has been proposed as the mechanism underlying within-area temporal dynamics at multiple stages of the ventral hierarchy (Ringach et al. [1997], Shapley et al. [2003], Carandini and Ringach [1997], Brincat and Connor [2006]). A local recurrent circuit operating on a characteristic timescale would leave a specific signature in the population activity: an enhancement in within-area predictivity at that timescale’s temporal offset, reflecting the additional structure that recurrent processing introduces beyond the shared feedforward drive. If such a circuit drives the representational complexity shifts, this enhancement should co-emerge with the shifts and plateau as they stabilize. In search of such a signal, we fit time-lagged ridge regression models that predicted an electrode’s response in a given time bin from same-area population activity in earlier time bins, varying the temporal offset between predictor and target.

Within-area time-lagged regression revealed a predictivity enhancement at a 30 ms temporal offset, present in all three areas in both monkeys. For each area in each monkey, predictivity generally decreased with increasing temporal separation, as expected (Fig. 6A). This otherwise monotonic decay was interrupted by a bump at 30 ms: activity 30 ms in the past predicted the current response better than activity 20 ms in the past. The non-monotonic profile indicates that population activity at this specific temporal offset carries additional predictive information beyond what a simple decay of shared stimulus drive would produce.

The 30 ms signal has the spatial and temporal properties expected of a local recurrent process. The regression weights at the 30 ms offset decayed progressively with inter-electrode distance before reaching a plateau at longer ranges (Fig. 6B). This short-range spatial profile was consistent across areas and monkeys and is compatible with the known spatial scale of intrinsic horizontal connections within cortical areas. The bump was absent or negative early in the response and grew progressively larger in later time bins across all three areas (Fig. 6C, left). Plotting the magnitude of the bump against the most-predictive model layer at each time bin revealed that the enhancement emerged alongside the representational complexity shifts and plateaued as the best layer stabilized (Fig. 6C, right).

The temporal correspondence, together with the spatially local structure of the underlying weights, identifies local recurrence as a candidate mechanism driving the complexity shifts, though we emphasize that the 30 ms signal does not by itself establish causality. Broadly distributed feedback operating on a similar timescale could in principle produce qualitatively similar within-area predictivity profiles. Further, the spatial decay of the 30 ms regression weights is consistent with the spatial scale of intrinsic horizontal connections, but cannot on its own distinguish a local circuit from a feedback projection that happens to act on a similar scale. Direct causal tests – for example, inactivating downstream areas while recording the shift in upstream populations – are required to settle this question.

### 2.7 Existing computational models do not capture the neural dynamics

Neural network models with temporal components offer a candidate explanation for the observed within-area complexity shifts. If a model’s dynamics matched those of the brain, providing it with additional temporal processing when predicting later neural time bins should reduce the apparent early-to-late shift in most-predictive layer for individual electrodes (Fig. 7A). We evaluated this hypothesis in two models with distinct temporal mechanisms. We employed CORnet-RT (Kubilius et al. [2018]), a convolutional network with local recurrence within each of its cortically mapped areas that was explicitly designed to produce dynamics in response to static images. We also tested V-JEPA 2 (Assran et al. [2025]), a self-supervised video transformer that could in principle produce shifts in static settings as a byproduct of learning to process dynamic stimuli. We applied the per-electrode shift analysis of Fig. 3 to both models, predicting the early time bin (35–45 ms) from each model’s initial features and the late time bin (185–195 ms) from features after additional temporal processing: additional biological unrolls for CORnet-RT, and an increased frame input for V-JEPA 2 (Fig. 7B, C). Here we report two additional unrolling steps for CORnet-RT and an additional 20 frames for V-JEPA 2.

Per-electrode shifts persisted under both dynamic mappings, suggesting that existing temporal models do not capture the same representational shifts as measured in the brain (Fig. 7). For both CORnet-RT (Fig. 7B) and V-JEPA 2 (Fig. 7C), electrodes in V4 and IT clustered above the diagonal in all conditions, indicating that the late time bin was best predicted by a deeper model layer than the early time bin even when the model was given additional temporal processing for the late bin. V1 followed the same pattern under V-JEPA 2. To quantify these shifts, we computed each electrode’s early-to-late layer shift (late best layer minus early best layer) and tested the resulting distribution against zero (one-sided t-test). Per-electrode shifts were significantly greater than zero in V4 and IT for both models and in V1 for V-JEPA 2 (all p *<* 0.001), but were not significantly different from zero in V1 under CORnet-RT (p = 0.215).

These results indicate that the within-area representational shifts are not reproduced by current state-of-the-art dynamic models. Expanding either model’s temporal processing does not reshape its features along the complexity axis we measured in the neural data. While CORnet-RT may capture some of the V1 shift, this result is hard to interpret given the model’s coarse layer-index resolution (only five layers), which limits sensitivity to the small shift expected in V1.

## 3 Discussion

Here, we show that within-area temporal dynamics across the ventral visual stream reflect a single shared computational motif: within each area, representations evolve along the same axis of representational complexity that organizes the visual hierarchy. That the same operation recurs across V1, V4, and IT, areas otherwise quite different in their selectivity, points to a shared underlying computational principle, and by extension a shared mechanism whose specific circuit-level implementation remains to be established. The shifts were broadly distributed across the recorded population rather than concentrated in a subpopulation and were dissociable from response timing within-area, characterizing the shift as a distributed area-wide property of each population. The shifts were functionally consequential, allowing more complex images to be decoded later in the response. We found evidence of a within-area predictivity enhancement at a 30 ms temporal offset, present in all three areas in both monkeys, consistent with a local recurrent circuit that may drive these shifts, motivating future work on underlying mechanism. Finally, we show that state-of-the-art dynamic models – including one with built-in local recurrence – did not reproduce the shifts measured in the data, suggesting a target for future modeling work.

The calibrated model hierarchy we employed puts within-area dynamics on the same axis as cross-area differences, unifying phenomena that prior approaches could not directly compare in a quantifiable way. Past characterizations measured within-area dynamics idiosyncratically between areas – orientation bandwidth in V1, multipart-shape selectivity in V4, global-to-fine identity coding in IT (Ringach et al. [1997], Brincat and Connor [2006], Sugase et al. [1999]). These axes are largely qualitative and do not generalize across the hierarchy. We instead quantified time-resolved deviations from a time-averaged feedforward baseline, supplied either by a brain-like deep neural network model or by another monkey’s actual brain. Both forms revealed the same motif: each area’s representations shifted over time, toward deeper model layers and higher cortical areas in the other monkey, with the latter grounding the result independently of any model. While previous work (Xiao et al. [2025]) suggests that the modelbased version of this finding applies within IT, we extend the theory across the visual hierarchy, with corroboration from the brain-as-baseline form.

Our results unify a collection of area-specific mechanistic hypotheses into a single proposal: that local recurrence may drive the observed complexity shifts. Local recurrence has been independently advanced as the mechanism underlying within-area dynamics in each of V1, V4, and IT (Ringach et al. [1997], Carandini and Ringach [1997], Shapley et al. [2003], Brincat and Connor [2006], Yau et al. [2013], Sugase et al. [1999], Matsumoto et al. [2005]). We observed a 30 ms within-area predictivity enhancement of consistent timescale and short-range spatial profile across all three areas – a signature whose form is consistent with local recurrent processing. This signature, present in all three areas, suggests local recurrence as a parsimonious candidate mechanism: a single circuit motif repeated at each stage rather than three distinct mechanisms producing similar dynamics.

Although our data are consistent with local circuitry contributing to the measured shifts, they do not rule out long-range feedback, especially in other regimes. Long-range feedback is well-documented to alter representations – for example, during figure-ground segregation (Lamme and Roelfsema [2000]), perceptual filling-in of occluded image regions in V1 of these same monkeys (Papale et al. [2023]), top-down attention and decision-making (Gilbert and Li [2013]), and behavioral tasks in which causal manipulations have confirmed its role (Kar and DiCarlo [2021], Kirchberger et al. [2021]). These effects are typically engaged during deliberate tasks or decision-making; the present recordings were collected during passive fixation.

Moreover, we attempted to quantify inter-area contributions via predictive modeling, but the interpretations were confounded. We predicted each electrode’s response from recent population history in different combinations of areas, with the intention to read the fit weights as a connectivity diagram (Supplementary Fig. 8). In practice, same-area history alone predicted responses as well as history from all areas combined, leaving little residual variance for cross-area predictors to explain. This saturation also obscured signatures of known feedforward drive. For example, V1 history did not boost V4 predictions above the same-area baseline, despite V1’s well-characterized projection to V4 – indicating that shared stimulus drive and within-area tuning correlations render the prediction problem too easy for fit weights to reflect underlying circuitry. Thus, while our autoregressive modeling approach failed to find evidence for long-range contributions to within-area dynamics, it also failed to recover the feedforward drive that we know is present, calling into question the validity of the approach.

Although we cannot directly resolve single-neuron dynamics because of the nature of multi-unit activity, prior single-unit work supports the single-neuron interpretation of these shifts. Specifically, MUA cannot distinguish two scenarios that would produce the same population-level shift: multiple neurons measured by a single electrode gradually shifting their tuning, or differently-tuned neurons being recruited at staggered latencies. However, single-unit recordings across V1, V4, and IT consistently show within-neuron tuning changes over the response (Ringach et al. [1997], Mazer et al. [2002], Brincat and Connor [2006], Sugase et al. [1999], Matsumoto et al. [2005]), supporting our interpretation that the shifts we measure reflect single-unit changes in tuning rather than a redistribution of which neurons contribute to the recorded signal.

The dynamics we characterize provide a direct target for models of the ventral visual stream, beyond second-order metrics. Current dynamic models, like CORnet-RT (Kubilius et al. [2018]) and recurrent CNNs (Nayebi et al. [2018, 2022]), have largely been validated against derived metrics, such as object solution time (Kar et al. [2019]). However, this type of test is not a first-order test of dynamic alignment. The within-area trajectories we report allow for a more direct test: whether a model’s internal representations actually align with neural representations at each time bin. By this stricter criterion, neither CORnet-RT nor V-JEPA 2 reproduced the trajectory, despite claims that CORnet-RT matches neural representational dynamics based on its ability to match object solution times. While some temporal models have been tested directly against neural data (e.g. Nayebi et al. [2022]), we suggest that the present data provide a richer target for evaluation due to the number of images and recorded units.

Ultimately, identifying why these dynamics emerge requires a goal-driven approach of comparing models optimized under different objectives. Our finding that harder images were decoded later is a functional correlation, not an explanation. Goal-driven modeling has produced normative accounts of static ventral stream representations by asking which architecture, training objective, training data, or learning rule produces brain-like features (Yamins et al. [2014], Yamins and DiCarlo [2016], Richards et al. [2019]). Analogous accounts for temporal dynamics remain open. However, our results suggest that the present neural data contain enough dynamic signal to constrain these goal-driven models and test mechanistic hypotheses.

## Acknowledgments

We are thankful to both the Stanford NeuroAI lab and Gardner lab for helpful feedback on the work, as well as Aran Nayebi, Ko Kar, Konrad Kording, and Matteo Carandini.

## Data and materials availability

Code and data to replicate the analyses described in the paper will be released publicly upon publication.

## 4 Methods

### 4.1 Data and processing

Multi-unit activity (MUA) data were obtained from the THINGS Ventral Stream Spiking Dataset (TVSD) (Papale et al. [2025]), which comprises large-scale electrophysiological recordings from two macaques (Monkeys F and N). Each animal was implanted with 16 64-channel Blackrock Utah arrays spanning V1, V4, and inferotemporal cortex (IT). Both monkeys had 8 V1 arrays; Monkey N additionally had 4 V4 arrays and 4 IT arrays, whereas Monkey F had 3 V4 arrays and 5 IT arrays. Each monkey viewed the same 22,348 unique images from the THINGS dataset (Hebart et al. [2019]), yielding 25,248 total trials each. 22,248 images were presented once (train set), and an additional 100 images were each presented 30 times (test set). Each image was presented for 200 ms while the monkeys maintained fixation. We relied on the established TVSD pipeline to extract and process data. Full acquisition and processing details are described in the original TVSD publication.

For analyses using time-averaged responses (Fig. 1), we used the single-trial normalized MUA responses provided in the TVSD. MUA was averaged over 100 ms windows aligned to image onset, with area-specific windows (V1: 25–125 ms; V4: 50–150 ms; IT: 75–175 ms). For time-resolved analyses (Figs. 2–7), we generated higher temporal resolution responses by applying the same extraction and averaging procedure used in the TVSD pipeline to compute normalized MUA averages over 10 ms bins. We extracted data for bin start times from 25 ms to 195 ms in 10 ms increments, though most analyses use 35-185 ms, as the 25 and 195 ms bins showed lower reliability.

### 4.2 Neural network feature extraction

We used DINOv2-Large as a fixed feature extractor for neural network-based analyses (Oquab et al. [2023]). We chose DINOv2 as the core model for our analyses as it had the highest neural predictivity of the candidate models we tested. DINOv2 is a self-supervised vision transformer trained with a DINO-style self-distillation objective to learn general-purpose visual representations from large-scale natural images (Caron et al. [2021]). The ViT-L/14 backbone (Dosovitskiy et al. [2020]) comprises 24 transformer blocks with a 1,024-dimensional embedding size and a 14 × 14 pixel patch size. We extracted features from the same RGB stimuli presented to the monkeys, resized to 256 × 256 and cropped to 224 × 224. Given a 14 × 14 patch size, each image is partitioned into a 16 × 16 grid of patches, yielding 256 patch tokens per image; we used the patch embeddings and CLS token, producing an array of shape (257, 1024) per image at each layer. For all analyses, we flattened the patch-token representation into a single feature vector per image per layer (257 × 1,024 features) without pooling, subsampling, or dimensionality reduction. For clarity in figures and summaries, we analyzed every other transformer block of the network, as adjacent blocks yielded highly similar predictivity profiles.

We include supplementary analyses with two additional networks, VGG19 (Simonyan and Zisserman [2014]) and CORnet-S (Kubilius et al. [2019]) to show that our results do not depend on the specific DINOv2 architecture. For VGG, we used the first convolutional layer, the five intermediate max pooling layers, the global pooling layer, and the readout head for a total of eight layers. For CORnet-S, we used the outputs of the V1, V2, V4, IT, and decoder blocks, for a total of five.

To assess whether a state-of-the-art dynamic vision model accounts for the within-area complexity shifts (Fig. 7), we additionally extracted features from V-JEPA 2 ViT-g (Assran et al. [2025]), a selfsupervised video transformer trained with a masked spatiotemporal prediction objective on large-scale natural video. Features were extracted from the encoder at seven evenly spaced layers (indices 0, 6, 12, 18, 24, 30, 36) using the same 224 × 224 RGB stimuli presented to the monkeys, in two input configurations. In the 1-frame configuration, the stimulus image was presented to the encoder as a single frame, yielding one set of output tokens per image per layer. In the 21-frame configuration, the stimulus image was replicated across 21 frames; because V-JEPA 2 tokenizes its input in space-time tubelets that each add a fixed number of tokens, the resulting output sequence grew with tubelet count. To match the token dimensionality of the 1-frame configuration, we retained only the tokens at the end of the 21-frame output sequence, corresponding to the most recently tokenized input region.

To assess whether a recurrent network with biologically plausible dynamics accounts for the within-area complexity shifts (Fig. 7), we additionally extracted features from CORnet-RT (Kubilius et al. [2018]), a shallow recurrent convolutional network that integrates local recurrent connections within each area via additive recurrence. Features were extracted from the output of each area (V1, V2, V4, IT, and the decoder) for a total of five layers. In the first-pass configuration, features were extracted after the first forward pass through each layer of the network. In the unrolled configuration, the network was run for two additional biological unrolls.

### 4.3 Linear mapping procedure

All linear mappings were implemented as ridge regression using RidgeCV in scikit-learn (sklearn). Ridge regression fits a linear model from predictors to target with an L2 penalty on the regression coefficients. We selected the regularization strength *α* ∈ [10^*−*10^, 10^10^] by leave-one-out cross-validation on the training set over log-spaced intervals. We determined *α* separately for each electrode. The same fitting procedure was used for both cross-animal mappings (neural responses as predictors) and model-to-brain mappings (network features as predictors). For time-resolved cross-animal ridge analyses (Supplementary Fig. 2B), we predicted time-resolved electrode responses in one animal from time-averaged responses in each area of the other animal, paralleling the asymmetric design used for cross-animal temporal RSA (see Representational similarity analysis).

Predictive mappings were fit for each electrode to maximize predictivity on the train set (minimize residual sum of squares with an L2 penalty), then predictivity was quantified using out-of-sample performance on the test set. Predictivity was computed either as the coefficient of determination (*r*^2^) or as the Pearson correlation (*r*) between predicted and observed responses, depending on the analysis. For evaluation on the test set, responses were averaged over the 30 repeats for each image, yielding one test response per image (100 test points per electrode) to reduce noise. Unless otherwise noted, population-level summaries report the median predictivity across all electrodes within an area. We did not apply noise-ceiling correction.

To assess whether DINOv2 reproduces the full pattern of cross-area predictivities (Fig. 1D), we determined each predictor area’s best-predicting DINOv2 layer (the layer yielding the highest median ridge predictivity for that area). We then used that layer’s features to predict responses in each of the three target areas. The resulting nine model predictivities were compared to the nine corresponding cross-animal predictivities via Pearson correlation.

### 4.4 Representational similarity analysis

We quantified representational alignment at the population level using representational similarity analysis (RSA) separately within each visual area via rsatoolbox (van den Bosch et al. [2025]). This evaluation was only performed on the test set. For each area and time bin, we computed a population response vector for each test image by averaging MUA across the 30 repeated presentations, yielding one population response per image. We then computed a neural representational similarity matrix (RSM) as the Pearson correlation between population response vectors for all pairs of test images, producing a 100 × 100 RSM for each area and time bin. We computed model RSMs analogously from DINOv2 features, and neural RSMs from corresponding or non-corresponding areas across the two animals. Alignment between any two RSMs was quantified as the Spearman correlation between their vectorized upper triangles, excluding the diagonal.

This procedure was applied in four configurations. For time-averaged analyses (Fig. 1C), we compared neural RSMs computed from 100 ms averaged responses to either model layer RSMs or neural RSMs from the other animal. For time-resolved analyses (Fig. 2A and Supplementary Fig. 1), we computed neural RSMs from 10 ms time bins and compared them to either model layer RSMs (Fig. 2A) or time-averaged neural RSMs from each area of the other animal (Supplementary Fig. 1). This design ensures that temporal shifts in alignment cannot be attributed to time-varying structure in the predictor RSM. For the cross-animal temporal RSA and ridge regression, tick marks indicating each time bin’s center of mass across predictor areas were computed by assigning ordinal indices to V1 (1), V4 (2), and IT (3) and applying a softmax.

### 4.5 Most-predictive layer estimation

We used two complementary measures of the most-predictive DINOv2 layer, depending on whether the analysis required a population-level or electrode-level summary. For population-level analyses (Figs. 2A, 5B, 6C), we computed a softmax-weighted center of mass over the alignment profile across DINOv2 layers. Alignment values – either Spearman correlations from RSA (Fig. 2) or Pearson correlations from ridge regression (Figs. 5B, 6C) – were passed through a softmax function with temperature 0.1, and the resulting weights were used to compute the weighted average of layer indices. The low temperature sharpens the weighting toward the peak of the profile while retaining a continuous, differentiable summary of the best-predicting network depth for a given area and time bin.

For electrode-level analyses, we determined the most-predictive DINOv2 layer for each electrode independently as the layer yielding the highest ridge regression predictivity (argmax). Area-level summaries (thick lines in Fig. 2B; curves in Fig. 4E) report the mean of these per-electrode argmaxes at each time bin. This approach preserves the distribution of best layers across individual electrodes. For analyses comparing early and late best layers (Fig. 3C, D), only electrodes with split-half reliability exceeding 0.2 in the earliest time bin (35–45 ms) were included, to ensure that early-time-bin layer assignments were based on meaningful signal. Additionally, only electrodes with an average split-half reliability *>* 0.2 across time bins were included. For the latency-versus-complexity analysis (Fig. 4B), each electrode’s ending best layer was computed as the mean of its per-time-bin argmax values over the last five time bins, rather than a single time bin, to reduce variation. Latencies (from Papale et al. [2025]) were log-transformed before regression against most-predictive ending layer.

### 4.6 Split-half reliability

We quantified the split-half reliability of each electrode by computing response consistency over stimulus repetitions in the test set. Split-half reliability was computed for both the 100 ms time-averaged responses and each of the 10 ms time-resolved response bins. For a given electrode and time bin, we randomly partitioned the 30 repeated presentations of each of the 100 test images into two halves, averaged responses within each half separately, and obtained two length-100 response vectors (one value per test image). Split-half reliability was defined as the Pearson correlation between these two vectors. We repeated this procedure 50 times with independent random splits and calculated the mean correlation as the reliability estimate for that electrode and time bin.

### 4.7 Embeddings and clustering

We visualized heterogeneity in both time-resolved representational complexity (most-predictive layer over time bins) and response timing (split-half reliability over time bins) by embedding electrode-wise temporal trajectories into two dimensions using t-SNE. The t-SNE procedure was identical for both embeddings and used a two-dimensional embedding initialized with PCA, perplexity 50, and an automatically determined learning rate. For reliability-based embeddings, we z-scored each electrode’s reliability timecourse across time bins (mean-subtracting and dividing by the across-bin standard deviation) prior to embedding so that the visualization emphasized differences in temporal profile rather than overall reliability magnitude.

To visualize how electrodes from different visual areas relate to one another in their temporal trajectories, we computed combined two-dimensional t-SNE embeddings across electrodes from V1, V4, and IT for both the most-predictive-layer trajectories (Supplementary Fig. 5A) and the reliability trajectories (Supplementary Fig. 5B), coloring each electrode by its visual area.

To facilitate comparisons of representational complexity across electrodes with distinct timing profiles within each area, we additionally grouped electrodes within each area using k-means clustering (sklearn) applied to the z-scored reliability timecourses (Fig. 4C-E). For this clustering analysis, t-SNE was computed separately for each of the three visual areas to visualize within-area structure. We used three clusters to capture coarse slow, medium, and fast timing profiles for analysis and visualization, without implying discrete physiological subpopulations. Cluster assignments were computed in the original reliability-timecourse feature space and then overlaid on the corresponding two-dimensional t-SNE embedding for display.

### 4.8 Image-level classification and solution-time analysis

To assess the functional consequences of temporal shifts in representational complexity, we trained logistic regression classifiers to decode image category from time-resolved neural population activity. We first pooled electrodes from both monkeys into a single population vector for each area (V1, V4, IT). We then trained classifiers separately for each area and each 10 ms time bin using MUA responses to the 1,200 training-set presentations (100 categories × 12 images per category) and evaluated on the 100 test-set images, with responses averaged across the 30 repeated presentations per image. A separate set of classifiers was trained on the time-averaged (100 ms window) responses for comparison. Classification was performed using logistic regression (scikit-learn LogisticRegression).

We defined an image as solved at the earliest time bin for which its correct category appeared in the top 5 classifier predictions for two consecutive time bins (the object solution time, or OST). Once solved, an image remained counted as solved for all subsequent time bins. Cumulative solution curves were computed as the number of test images solved at or before each time bin. Empirically, we found this 2-bin top-5 criterion to be a good tradeoff between flexibility and accuracy for the classifier, given the constraints of the noisy neural data and the limited training examples.

To compare neural and network solution times at the level of individual images, we independently determined each image’s solution time in the neural data (as described above) and solution layer in DINOv2. For the network, a logistic regression classifier was trained at each DINOv2 layer to predict image category from layer features, using the same 1,200 training images. An image was defined as solved at the earliest layer for which the correct category was the top-1 prediction for two consecutive layers. The more stringent top-1 criterion was used for the network because network representations are noiseless, and the classifier performance was much higher. To relate classification performance to representational complexity (Fig. 5B), we plotted cumulative images solved at each time bin against the most-predictive DINOv2 layer for that area and time bin, computed as the population-level softmax-weighted center of mass (see Most-predictive layer estimation).

### 4.9 Within-area time-bin mapping and weight analysis

To characterize temporal dependencies within each visual area, we fit ridge regression models that mapped population activity in one time bin to population activity in a later time bin within the same area. For each area, the predictor set consisted of all electrodes recorded in that area, including the target electrode itself. We considered all 10 ms time bins from 25 ms through 195 ms and fit mappings only in the forward direction (predictor time earlier than target time). Models were trained on the training image set and evaluated out of sample on the repeated-image test set using responses averaged across the 30 repeats for each test image, as described above. The regularization strength *α* was selected by leave-one-out cross-validation on the training set over the same *α* grid used for the other linear mappings. Mappings were done separately in each monkey.

To relate the fitted mappings to array-scale spatial organization, we analyzed how regression weights varied with inter-electrode distance for the 30 ms offset mapping, corresponding to the offset exhibiting local enhancement in predictability. This analysis was performed separately for each predicted time bin. Because electrode positions are known only within individual Utah arrays, the analysis was restricted to within-array electrode pairs. Electrodes within an array are spaced 0.4 mm apart, allowing interelectrode distances to be computed from array grid coordinates. For each distance, we summarized weight magnitude as the mean absolute regression coefficient across all electrode pairs separated by that distance, averaged across arrays within each monkey.

To track how the non-monotonic predictivity enhancement at the 30 ms offset (Fig. 6A) developed over the course of the response, we computed the magnitude of the predictivity bump for each predicted time bin and area (Fig. 6C). Bump magnitude was defined as the difference in out-of-sample predictivity (Pearson correlation) between the 30 ms offset and the adjacent 20 ms offset. A positive value indicates that same-area population activity 30 ms in the past predicted the current response better than activity 20 ms in the past, beyond what a monotonic decay from shared stimulus drive would produce. Values were averaged across both monkeys. To relate the emergence of this enhancement to the representational complexity shifts (Fig. 6C, right), we plotted bump magnitude against the most-predictive DINOv2 layer at each time bin, computed as the softmax-weighted center of mass of the population-level ridge regression predictivity profile across layers (temperature = 0.1; see Most-predictive layer estimation).

## 5 Supplementary information

### 5.1 Cross-animal temporal RSA

**Figure S1:**
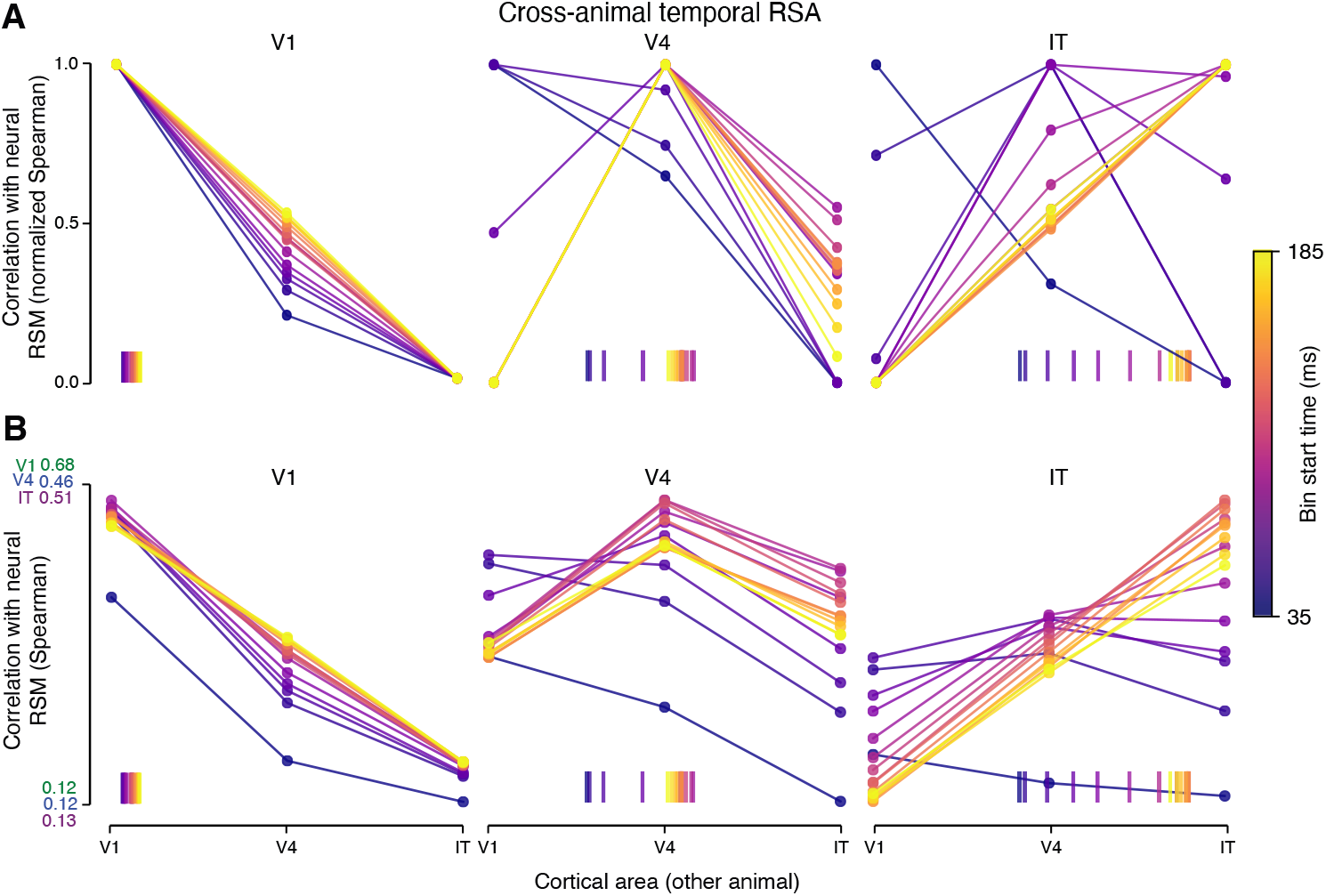
Cross-animal temporal RSA confirms the same complexity shifts as model-brain temporal RSA, without reference to an artificial network. For each time bin and target cortical area in one monkey (panel labels above each plot), we computed the Spearman correlation between the time-resolved neural RSM and the time-averaged RSM (100 ms window) of each area in the other monkey (predictor area, x-axis). Points are averages across both monkeys; color denotes time bin (35–185 ms post-stimulus). (A) Correlations normalized to span 0–1 within each time bin to isolate changes in the depth profile from changes in overall magnitude driven by time-varying reliability. (B) Same data without per-time-bin normalization. Tick marks below each plot indicate the softmax-weighted center of mass across predictor areas at each time bin (V1=1, V4=2, IT=3). In all three areas, representational geometry shifts over time toward higher cortical areas in the other monkey – for example, V1 representations become progressively more similar to the other monkey’s V4 over the course of the response.

### 5.2 Shifting ridge predictivity

**Figure S2:**
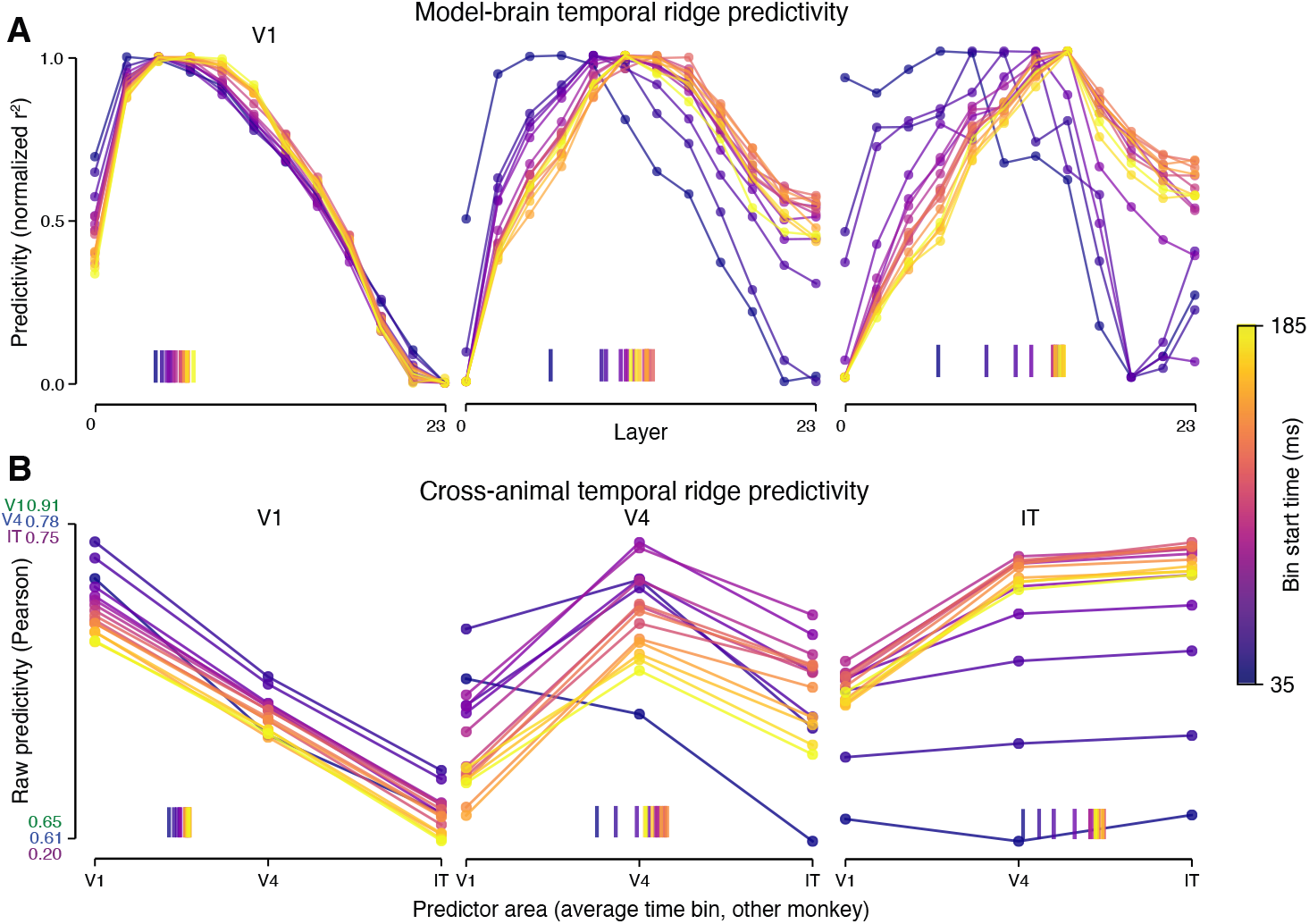
Time-resolved ridge regression recovers the same complexity shifts as RSA. (A) Model-brain temporal ridge predictivity. For each time bin (color) and cortical area, the mean out-of-sample ridge regression predictivity (r^2^) between DINOv2 layer features and neural responses across electrodes is shown, normalized to span 0–1 within each time bin. Tick marks indicate the softmax-weighted center of mass across layers for each time bin. (B) Cross-animal temporal ridge predictivity. For each time bin and cortical area in one monkey, raw Pearson predictivity from time-averaged responses (100 ms window) in each area of the other monkey to the time-resolved target responses is shown. Points are means across both monkeys. Tick marks indicate the center of mass (softmax) across predictor areas. Conventions for (A) as in Fig. 2A; conventions for (B) as in Supplementary Fig. 1.

### 5.3 Other models show the same shifting effect

**Figure S3:**
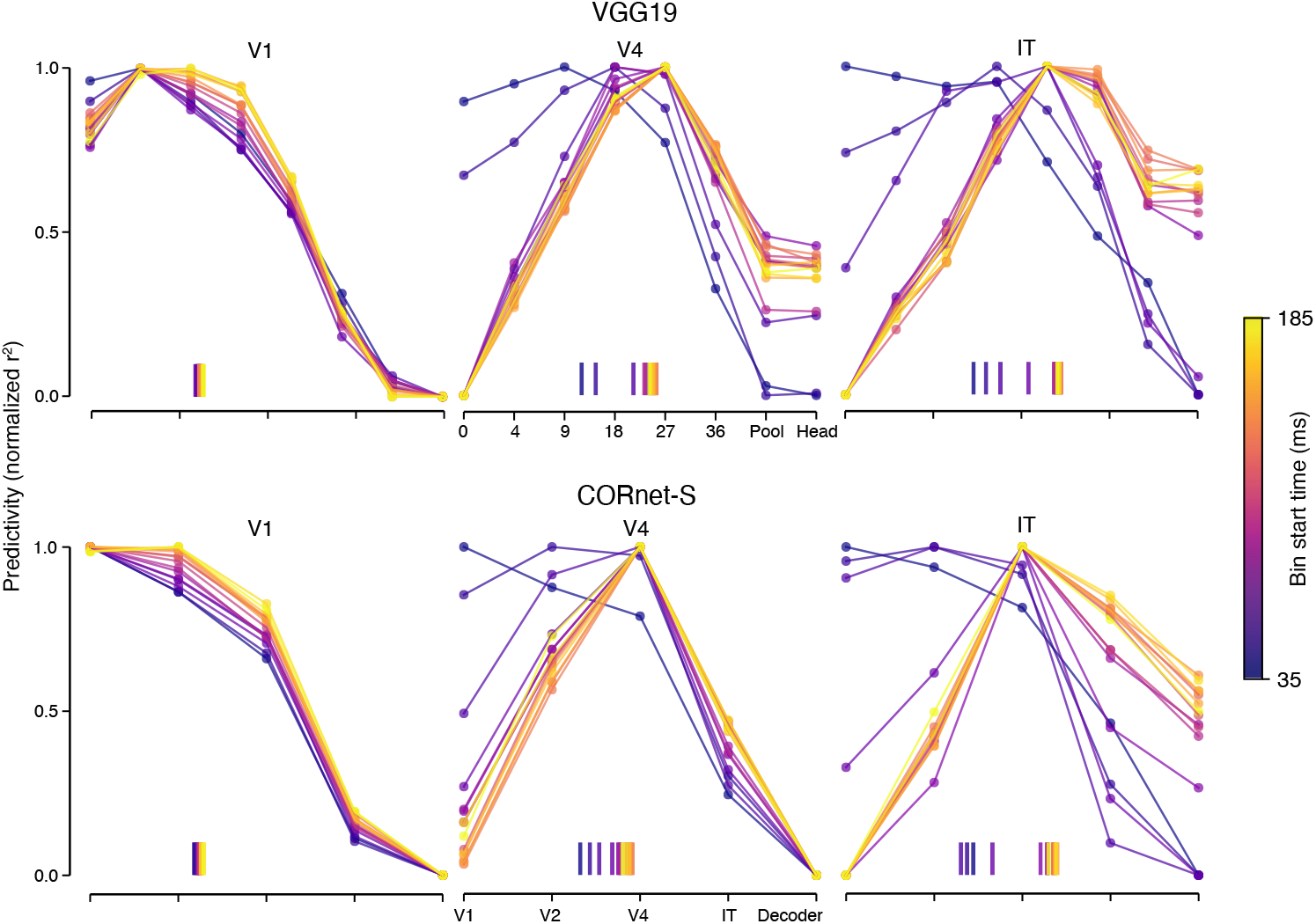
Temporal complexity shifts replicate with VGG19 and CORnet-S. Model-brain temporal ridge predictivity for VGG19 (top) and CORnet-S (bottom). For each time bin (color) and cortical area, normalized predictivity (r^2^) is shown across successive layers of each network. Tick marks indicate the softmax-weighted center of mass across layers. In both networks, the peak of the predictivity profile shifts toward deeper layers over the course of the response in all three areas, matching the pattern observed with DINOv2.

### 5.4 Unnormalized shifts

**Figure S4:**
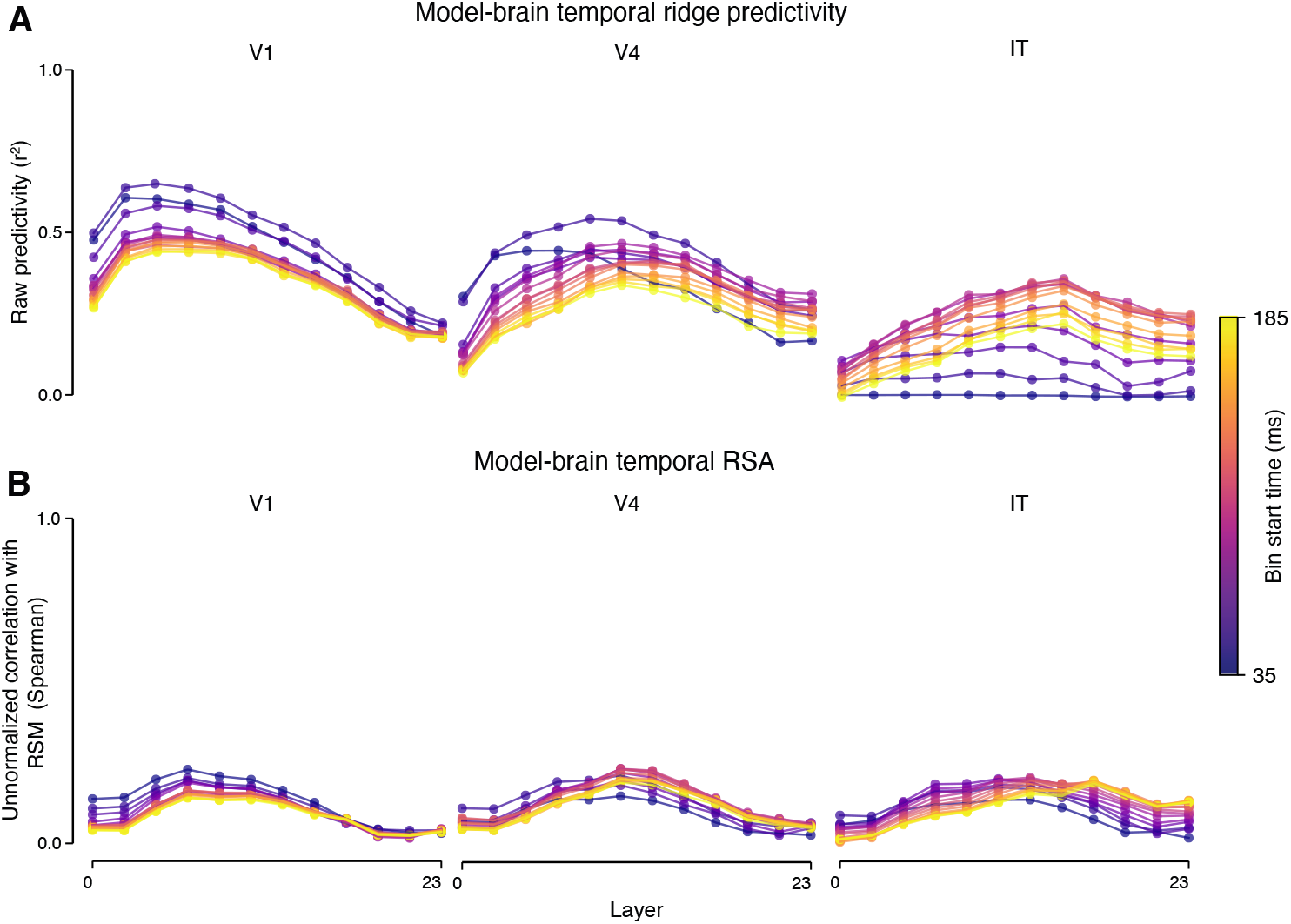
Complexity shifts are preserved without normalization. (A) Model-brain temporal ridge predictivity and (B) model-brain temporal RSA, shown without per-time-bin normalization. Conventions as in Fig. 2A. Overall magnitudes vary across time bins due to differences in signal reliability, but the location of the peak along the layer axis – and thus the softmax-weighted center of mass used throughout the paper – is unaffected by normalization.

### 5.5 Combined-area t-SNE embeddings

**Figure S5:**
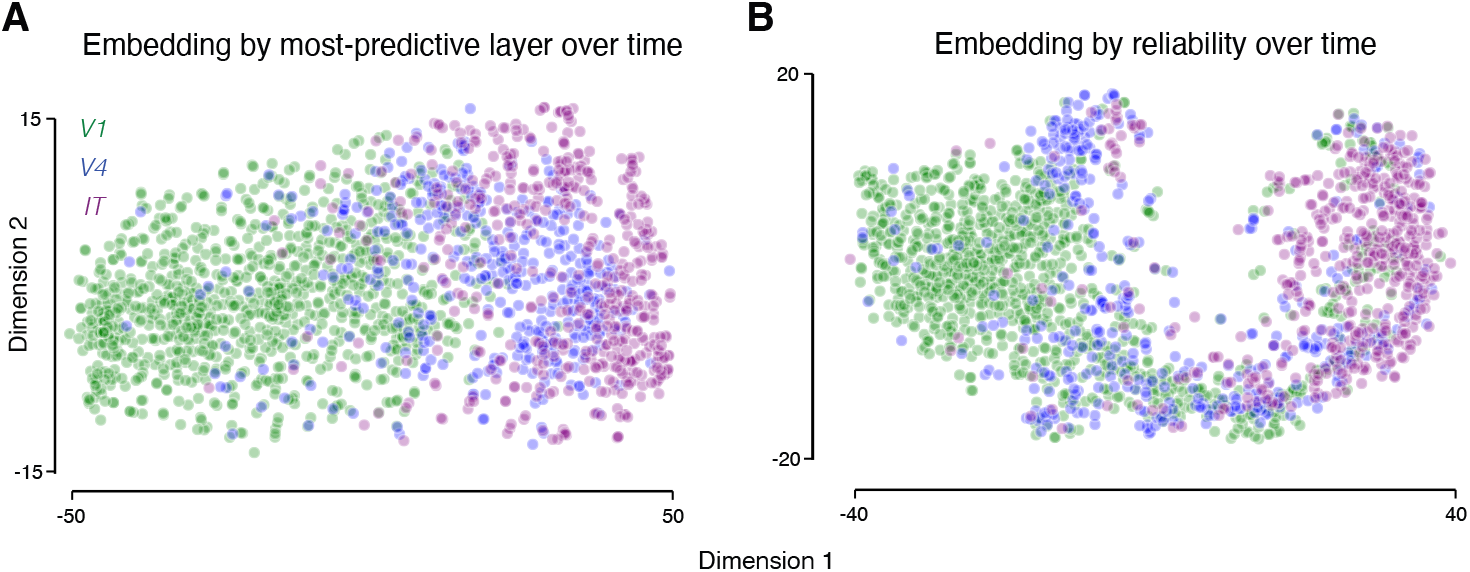
t-SNE embeddings of electrode-wise temporal trajectories reveal partially overlapping but distinguishable clusters by visual area. (A) Two-dimensional t-SNE embedding of each electrode’s most-predictive model layer over time, computed across all electrodes from V1, V4, and IT in a single embedding space. Each point is one electrode; color denotes its visual area (green, V1; blue, V4; purple, IT). (B) Two-dimensional t-SNE embedding of each electrode’s z-scored split-half reliability timecourse, also computed across all three areas combined. Conventions as in (A). Both embeddings reveal partially overlapping but distinguishable clusters corresponding to V1, V4, and IT, indicating that an electrode’s temporal complexity trajectory and its temporal reliability profile each carry information about its hierarchical position.

### 5.6 Increasing reliability does not account for more solved images

**Figure S6:**
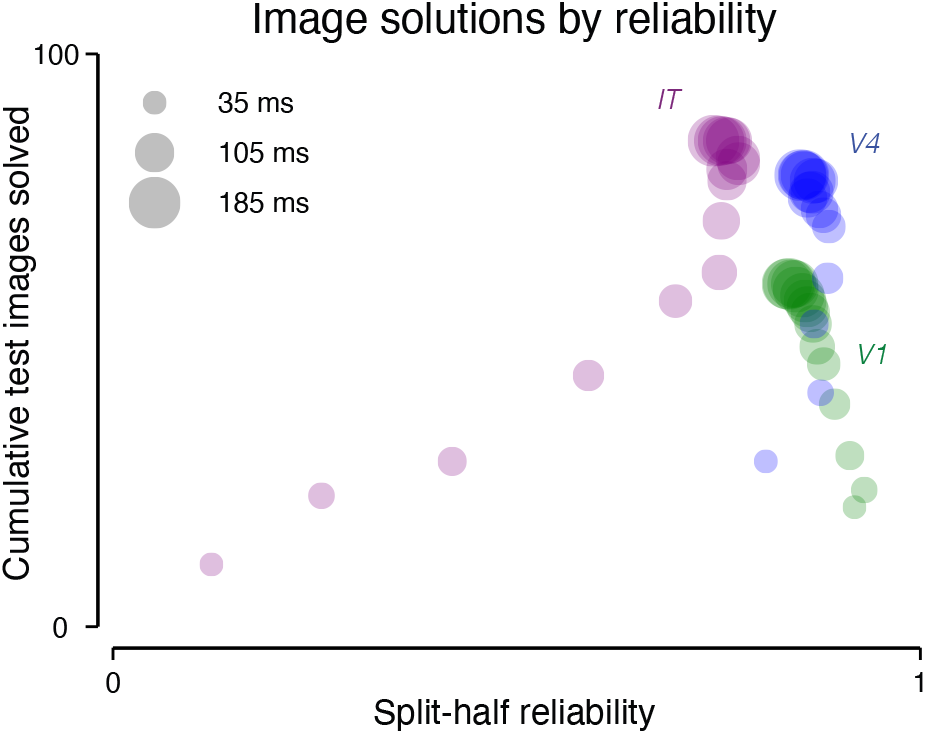
Classification improvements are only partially explained by increases in signal reliability. Cumulative number of test images solved by logistic regression classifiers trained on neural data from each area (color: green, V1; blue, V4; purple, IT) plotted against mean split-half reliability at each time bin. Bubble size indicates time bin (larger = later).

### 5.7 Cross-animal OST correspondence

**Figure S7:**
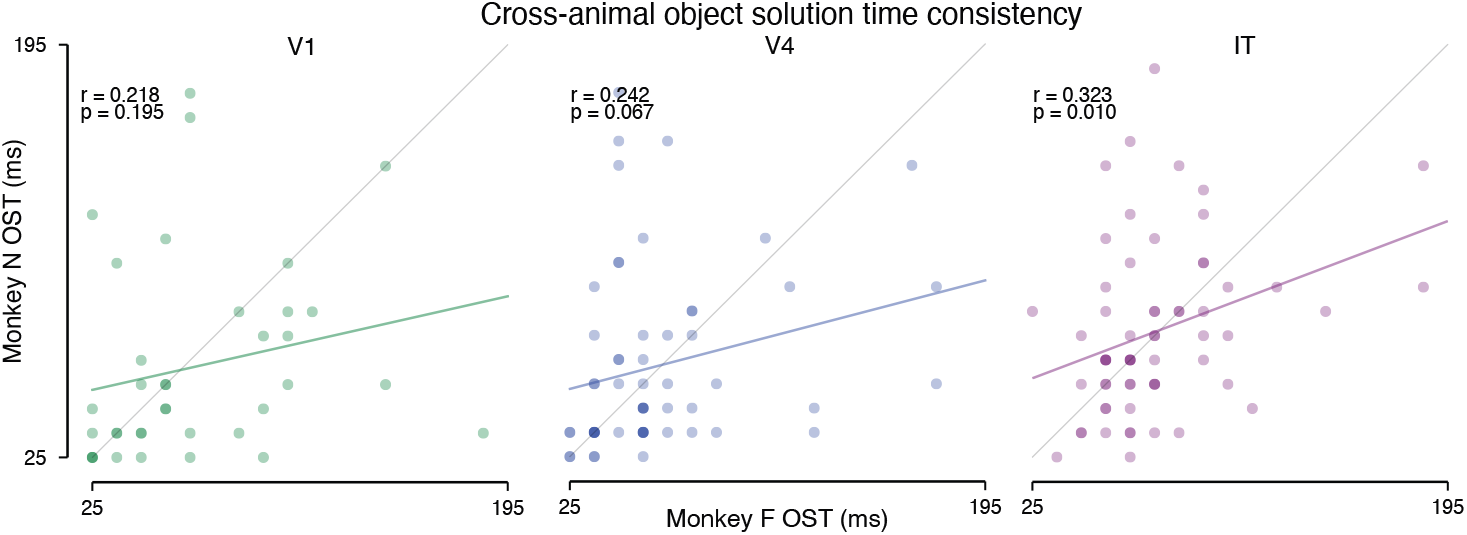
The two animals show weak but correlated object solution times. For each area, each point shows a single image’s object solution time in Monkey N (y-axis) plotted against its solution time in Monkey F (x-axis). Images are considered solved when the true class is in the top 5 predictions for two consecutive time bins.

### 5.8 Autoregressive modeling of neural data

**Figure S8:**
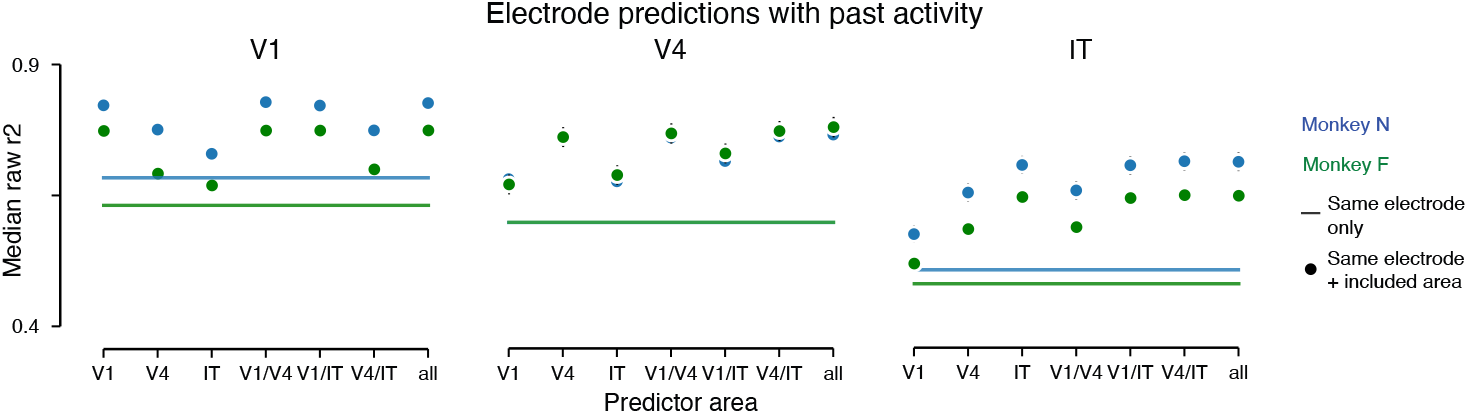
Inter-area autoregressive modeling is confounded. For each target electrode, ridge regression models were fit to predict its response in time bin t from past population activity. Panels show results for different target areas. Horizontal lines show mean predictivity across electrodes when using only the target electrode’s own response from the immediately preceding 10 ms time bin (t - 10 ms). Dots show mean predictivity when the target electrode’s own t - 10 ms response is supplemented with the past 50 ms of population activity (5 consecutive time bins) from electrodes in the area or combination of areas indicated on the x-axis. Predictivity is averaged across all target electrodes and all predicted time bins from 75 ms onward – the earliest bin for which 50 ms of preceding population activity is available. Colors denote monkey (blue, Monkey N; green, Monkey F).

To examine whether long-range inputs from other areas contribute to within-area dynamics beyond local processing, we fit ridge regression models that predicted each electrode’s response from past population activity, varying which areas contributed to the predictor set. The baseline predictor was the target electrode’s own response in the immediately preceding 10 ms time bin (t - 10 ms). We then expanded the predictor set to include the past 50 ms of population activity from electrodes in either a single area (V1, V4, or IT), a pair of areas, or all three areas combined, while always retaining the target electrode’s own t - 10 ms response. Predictivity was evaluated on held-out test images and averaged across all target electrodes and all predicted time bins from 75 ms onward, the earliest bin for which 50 ms of preceding population activity was available.

Across all three target areas and both monkeys, predictivity was largely saturated by same-area history (Supplementary Fig. 8). Adding same-area population history to the autoregressive baseline produced a substantial boost over the same-electrode case, while adding population history from other areas on top of same-area history yielded little further improvement. Including only non-target areas (e.g., V4 alone or IT alone for V1 targets) produced smaller gains over the baseline than including the target area’s own history.

This analysis did not, however, recover signatures of known feedforward connectivity: V1 population history did not measurably boost V4 predictivity above the same-area baseline, despite V1’s well-characterized feedforward projection to V4. The most plausible explanation is that shared stimulus drive across areas and within-area tuning correlations render same-area history a near-sufficient statistic for current activity regardless of the underlying circuit-level mechanism, obscuring both feedforward and feedback inter-area contributions. We therefore treat the analysis as uninformative about the relative roles of long-range and local processing.

## References

Daniel Anthes, Sushrut Thorat, Anna Mitola, Paolo Papale, Peter Kö nig and Tim C Kietzmann. The illusory simplicity of the feedforward pass: evidence for the dynamical nature of stimulus encoding along the primate ventral stream. arXiv, 2026.

Mido Assran, Adrien Bardes, David Fan, Quentin Garrido, Russell Howes, Mojtaba, Komeili, Matthew Muckley, Ammar Rizvi, Claire Roberts, Koustuv Sinha, Artem Zholus, Sergio Arnaud, Abha Gejji, Ada Martin, Francois Robert Hogan, Daniel Dugas, Piotr Bojanowski, Vasil Khalidov, Patrick Labatut, Francisco Massa, Marc Szafraniec, Kapil Krishnakumar, Yong Li, Xiaodong Ma, Sarath Chandar, Franziska Meier, Yann LeCun, Michael Rabbat, and Nicolas Ballas. V-JEPA 2: Self-supervised video models enable understanding, prediction and planning. arXiv, 2025. doi: 10.48550/arxiv.2506.09985.

C. E. Bredfeldt and D. L. Ringach. Dynamics of spatial frequency tuning in macaque v1. The Journal of Neuroscience, 22(5):1976–1984, 2002. ISSN 0270-6474. doi: 10.1523/jneurosci.22-05-01976.2002.

Scott L. Brincat and Charles E. Connor. Dynamic shape synthesis in posterior inferotemporal cortex. Neuron, 49(1):17–24, 2006. ISSN 0896-6273. doi: 10.1016/j.neuron.2005.11.026.

Santiago A. Cadena, George H. Denfield, Edgar Y. Walker, Leon A. Gatys, Andreas S. Tolias, Matthias Bethge, and Alexander S. Ecker. Deep convolutional models improve predictions of macaque v1 responses to natural images. PLoS Computational Biology, 15(4):e1006897, 2019. ISSN 1553-734X. doi: 10.1371/journal.pcbi.1006897.

Matteo Carandini and David J. Heeger. Normalization as a canonical neural computation. Nature Reviews Neuroscience, 13(1):51–62, 2012. ISSN 1471-003X. doi: 10.1038/nrn3136.

Matteo Carandini and Dario L. Ringach. Predictions of a recurrent model of orientation selectivity. Vision Research, 37(21):3061–3071, 1997. ISSN 0042-6989. doi: 10.1016/s0042-6989(97)00100-4.

Matteo Carandini, Jonathan B. Demb, Valerio Mante, David J. Tolhurst, Yang Dan, Bruno A. Olshausen, Jack L. Gallant, and Nicole C. Rust. Do we know what the early visual system does? The Journal of Neuroscience, 25(46):10577–10597, 2005. ISSN 0270-6474. doi: 10.1523/jneurosci.3726-05.2005.

Mathilde Caron, Hugo Touvron, Ishan Misra, Hervé Jégou, Julien Mairal, Piotr Bojanowski, and Armand Joulin. Emerging properties in self-supervised vision transformers. arXiv, 2021.

R Desimone, TD Albright, CG Gross, and C Bruce. Stimulus-selective properties of inferior temporal neurons in the macaque. The Journal of Neuroscience, 4(8):2051–2062, 1984. ISSN 0270-6474. doi: 10.1523/jneurosci.04-08-02051.1984.

James J. DiCarlo and David D. Cox. Untangling invariant object recognition. Trends in Cognitive Sciences, 11(8):333–341, 2007. ISSN 1364-6613. doi: 10.1016/j.tics.2007.06.010.

James J. DiCarlo, Davide Zoccolan, and Nicole C. Rust. How does the brain solve visual object recognition? Neuron, 73(3):415–434, 2012. ISSN 0896-6273. doi: 10.1016/j.neuron.2012.01.010.

Alexey Dosovitskiy, Lucas Beyer, Alexander Kolesnikov, Dirk Weissenborn, Xiaohua Zhai, Thomas Unterthiner, Mostafa Dehghani, Matthias Minderer, Georg Heigold, Sylvain Gelly, Jakob Uszkoreit, and Neil Houlsby. An image is worth 16 × 16 words: Transformers for image recognition at scale. arXiv, 2020. doi: 10.48550/arxiv.2010.11929.

Daniel J. Felleman and David C. Van Essen. Distributed hierarchical processing in the primate cerebral cortex. Cerebral Cortex, 1(1):1–47, 1991. ISSN 1047-3211. doi: 10.1093/cercor/1.1.1-a.

Jeremy Freeman, Corey M Ziemba, David J Heeger, Eero P Simoncelli, and J Anthony Movshon. A functional and perceptual signature of the second visual area in primates. Nature Neuroscience, 16 (7):974–981, 2013. ISSN 1097-6256. doi: 10.1038/nn.3402.

Jack L. Gallant, Jochen Braun, and David C. Van Essen. Selectivity for polar, hyperbolic, and cartesian gratings in macaque visual cortex. Science, 259(5091):100–103, 1993. ISSN 0036-8075. doi: 10.1126/science.8418487.

Charles D. Gilbert and Wu Li. Top-down influences on visual processing. Nature Reviews Neuroscience, 14(5):350–363, 2013. ISSN 1471-003X. doi: 10.1038/nrn3476.

Melvyn A. Goodale and A.David Milner. Separate visual pathways for perception and action. Trends in Neurosciences, 15(1):20–25, 1992. ISSN 0166-2236. doi: 10.1016/0166-2236(92)90344-8.

C G Gross, C E Rocha-Miranda, and D B Bender. Visual properties of neurons in inferotemporal cortex of the macaque. Journal of Neurophysiology, 35(1):96–111, 1972. ISSN 0022-3077. doi: 10.1152/jn.1972.35.1.96.

Umut Gü çlü and Marcel A. J. van Gerven. Deep neural networks reveal a gradient in the complexity of neural representations across the ventral stream. The Journal of Neuroscience, 35(27):10005–10014, 2015. ISSN 0270-6474. doi: 10.1523/jneurosci.5023-14.2015.

James V. Haxby, M. Ida Gobbini, Maura L. Furey, Alumit Ishai, Jennifer L. Schouten, and Pietro Pietrini. Distributed and overlapping representations of faces and objects in ventral temporal cortex. Science, 293(5539):2425–2430, 2001. ISSN 0036-8075. doi: 10.1126/science.1063736.

Martin N. Hebart, Adam H. Dickter, Alexis Kidder, Wan Y. Kwok, Anna Corriveau, Caitlin Van Wicklin, and Chris I. Baker. THINGS: A database of 1,854 object concepts and more than 26,000 naturalistic object images. PLoS ONE, 14(10):e0223792, 2019. doi: 10.1371/journal.pone.0223792.

D. H. Hubel and T. N. Wiesel. Receptive fields of single neurones in the cat’s striate cortex. The Journal of Physiology, 148(3):574–591, 1959. ISSN 0022-3751. doi: 10.1113/jphysiol.1959.sp006308.

D. H. Hubel and T. N. Wiesel. Receptive fields, binocular interaction and functional architecture in the cat’s visual cortex. The Journal of Physiology, 160(1):106–154, 1962. ISSN 0022-3751. doi: 10.1113/jphysiol.1962.sp006837.

D. H. Hubel and T. N. Wiesel. Receptive fields and functional architecture of monkey striate cortex. The Journal of Physiology, 195(1):215–243, 1968. ISSN 0022-3751. doi: 10.1113/jphysiol.1968.sp008455.

Chou P. Hung, Gabriel Kreiman, Tomaso Poggio, and James J. DiCarlo. Fast readout of object identity from macaque inferior temporal cortex. Science, 310(5749):863–866, 2005. ISSN 0036-8075. doi: 10.1126/science.1117593.

M. Ito, H. Tamura, I. Fujita, and K. Tanaka. Size and position invariance of neuronal responses in monkey inferotemporal cortex. Journal of Neurophysiology, 73(1):218–226, 1995. ISSN 0022-3077. doi: 10.1152/jn.1995.73.1.218.

Akshay V. Jagadeesh and Justin L. Gardner. Texture-like representation of objects in human visual cortex. Proceedings of the National Academy of Sciences, 119(17):e2115302119, 2022. ISSN 0027-8424. doi: 10.1073/pnas.2115302119.

Kohitij Kar and James J. DiCarlo. Fast recurrent processing via ventrolateral prefrontal cortex is needed by the primate ventral stream for robust core visual object recognition. Neuron, 109(1):164–176.e5, 2021. ISSN 0896-6273. doi: 10.1016/j.neuron.2020.09.035.

Kohitij Kar, Jonas Kubilius, Kailyn Schmidt, Elias B. Issa, and James J. DiCarlo. Evidence that recurrent circuits are critical to the ventral stream’s execution of core object recognition behavior. Nature Neuroscience, 22(6):974–983, 2019. ISSN 1097-6256. doi: 10.1038/s41593-019-0392-5.

Seyed-Mahdi Khaligh-Razavi and Nikolaus Kriegeskorte. Deep supervised, but not unsupervised, models may explain IT cortical representation. PLoS Computational Biology, 10(11):e1003915, 2014. ISSN 1553-734X. doi: 10.1371/journal.pcbi.1003915.

Tim C. Kietzmann, Courtney J. Spoerer, Lynn K. A. Sö rensen Radoslaw M. Cichy, Olaf Hauk, and Nikolaus Kriegeskorte. Recurrence is required to capture the representational dynamics of the human visual system. Proceedings of the National Academy of Sciences, 116(43):21854–21863, 2019. ISSN 0027-8424. doi: 10.1073/pnas.1905544116.

Lisa Kirchberger, Sreedeep Mukherjee, Ulf H. Schnabel, Enny H. van Beest, Areg Barsegyan, Christiaan N. Levelt, J. Alexander Heimel, Jeannette A. M. Lorteije, Chris van der Togt, Matthew W. Self, and Pieter R. Roelfsema. The essential role of recurrent processing for figure-ground perception in mice. Science Advances, 7(27):eabe1833, 2021. doi: 10.1126/sciadv.abe1833.

Gabriel Kreiman and Thomas Serre. Beyond the feedforward sweep: feedback computations in the visual cortex. Annals of the New York Academy of Sciences, 1464(1):222–241, 2020. ISSN 0077-8923. doi: 10.1111/nyas.14320.

Nikolaus Kriegeskorte, Marieke Mur, and Peter A Bandettini. Representational similarity analysis - connecting the branches of systems neuroscience. Frontiers in Systems Neuroscience, 2:4, 2008. ISSN 1662-5137. doi: 10.3389/neuro.06.004.2008.

Jonas Kubilius, Martin Schrimpf, Aran Nayebi, Daniel Bear, Daniel L. K. Yamins, and James J. DiCarlo. CORnet: Modeling the neural mechanisms of core object recognition. bioRxiv, page 408385, 2018. doi: 10.1101/408385.

Jonas Kubilius, Martin Schrimpf, Kohitij Kar, Ha Hong, Najib J Majaj, Rishi Rajalingham, Elias B Issa, Pouya Bashivan, Jonathan Prescott-Roy, Kailyn Schmidt, Aran Nayebi, Daniel Bear, Daniel L K Yamins, and James J DiCarlo. Brain-like object recognition with high-performing shallow recurrent ANNs. arXiv, 2019. doi: 10.48550/arxiv.1909.06161.

Victor A.F. Lamme and Pieter R. Roelfsema. The distinct modes of vision offered by feedforward and recurrent processing. Trends in Neurosciences, 23(11):571–579, 2000. ISSN 0166-2236. doi: 10.1016/s0166-2236(00)01657-x.

N K Logothetis and D L Sheinberg. Visual object recognition. Annual Review of Neuroscience, 19(1): 577–621, 1996. ISSN 0147-006x. doi: 10.1146/annurev.ne.19.030196.003045.

Narihisa Matsumoto, Masato Okada, Yasuko Sugase-Miyamoto, Shigeru Yamane, and Kenji Kawano. Population dynamics of face-responsive neurons in the inferior temporal cortex. Cerebral Cortex, 15 (8):1103–1112, 2005. ISSN 1047-3211. doi: 10.1093/cercor/bhh209.

James A. Mazer, William E. Vinje, Josh McDermott, Peter H. Schiller, and Jack L. Gallant. Spatial frequency and orientation tuning dynamics in area v1. Proceedings of the National Academy of Sciences, 99(3):1645–1650, 2002. ISSN 0027-8424. doi: 10.1073/pnas.022638499.

Aran Nayebi, Daniel Bear, Jonas Kubilius, Kohitij Kar, Surya Ganguli, David Sussillo, James J DiCarlo, and Daniel L K Yamins. Task-driven convolutional recurrent models of the visual system. arXiv, 2018. doi: 10.48550/arxiv.1807.00053.

Aran Nayebi, Javier Sagastuy-Brena, Daniel M. Bear, Kohitij Kar, Jonas Kubilius, Surya Ganguli, David Sussillo, James J. DiCarlo, and Daniel L. K. Yamins. Recurrent connections in the primate ventral visual stream mediate a trade-off between task performance and network size during core object recognition. Neural Computation, 34(8):1652–1675, 2022. ISSN 0899-7667. doi: 10.1162/necoa01506.

Bruno A. Olshausen and David J. Field. Emergence of simple-cell receptive field properties by learning a sparse code for natural images. Nature, 381(6583):607–609, 1996. ISSN 0028-0836. doi: 10.1038/381607a0.

Maxime Oquab, Timothée Darcet, Théo Moutakanni, Huy Vo, Marc Szafraniec, Vasil Khalidov, Pierre Fernandez, Daniel Haziza, Francisco Massa, Alaaeldin El-Nouby, Mahmoud Assran, Nicolas Ballas, Wojciech Galuba, Russell Howes, Po-Yao Huang, Shang-Wen Li, Ishan Misra, Michael Rabbat, Vasu Sharma, Gabriel Synnaeve, Hu Xu, Hervé Jegou Julien Mairal, Patrick Labatut, Armand Joulin, and Piotr Bojanowski. DINOv2: Learning robust visual features without supervision. arXiv, 2023. doi: 10.48550/arxiv.2304.07193.

Paolo Papale, Feng Wang, A. Tyler Morgan, Xing Chen, Amparo Gilhuis, Lucy S. Petro, Lars Muckli, Pieter R. Roelfsema, and Matthew W. Self. The representation of occluded image regions in area v1 of monkeys and humans. Current Biology, 33(18):3865–3871.e3, 2023. ISSN 0960-9822. doi: 10.1016/j.cub.2023.08.010.

Paolo Papale, Feng Wang, Matthew W. Self, and Pieter R. Roelfsema. An extensive dataset of spiking activity to reveal the syntax of the ventral stream. Neuron, 2025. ISSN 0896-6273. doi: 10.1016/j.neuron.2024.12.003.

Anitha Pasupathy and Charles E. Connor. Responses to contour features in macaque area v4. Journal of Neurophysiology, 82(5):2490–2502, 1999. ISSN 0022-3077. doi: 10.1152/jn.1999.82.5.2490.

Anitha Pasupathy and Charles E. Connor. Shape representation in area v4: Position-specific tuning for boundary conformation. Journal of Neurophysiology, 86(5):2505–2519, 2001. ISSN 0022-3077. doi: 10.1152/jn.2001.86.5.2505.

Blake A. Richards, Timothy P. Lillicrap, Philippe Beaudoin, Yoshua Bengio, Rafal Bogacz, Amelia Christensen, Claudia Clopath, Rui Ponte Costa, Archy de Berker, Surya Ganguli, Colleen J. Gillon, Danijar Hafner, Adam Kepecs, Nikolaus Kriegeskorte, Peter Latham, Grace W. Lindsay, Kenneth D. Miller, Richard Naud, Christopher C. Pack, Panayiota Poirazi, Pieter Roelfsema, João Sacramento, Andrew Saxe, Benjamin Scellier, Anna C. Schapiro, Walter Senn, Greg Wayne, Daniel Yamins, Friedemann Zenke, Joel Zylberberg, Denis Therien, and Konrad P. Kording. A deep learning framework for neuroscience. Nature Neuroscience, 22(11):1761–1770, 2019. ISSN 1097-6256. doi: 10.1038/s41593-019-0520-2.

Maximilian Riesenhuber and Tomaso Poggio. Hierarchical models of object recognition in cortex. Nature Neuroscience, 2(11):1019–1025, 1999. ISSN 1097-6256. doi: 10.1038/14819.

Dario L. Ringach, Michael J Hawken, and Robert Shapley. Dynamics of orientation tuning in macaque primary visual cortex. Nature, 387(6630):281–284, 1997. ISSN 0028-0836. doi: 10.1038/387281a0.

Nicole C. Rust and James J. DiCarlo. Selectivity and tolerance (“invariance”) both increase as visual information propagates from cortical area v4 to IT. The Journal of Neuroscience, 30(39):12978–12995, 2010. ISSN 0270-6474. doi: 10.1523/jneurosci.0179-10.2010.

Robert Shapley, Michael Hawken, and Dario L. Ringach. Dynamics of orientation selectivity in the primary visual cortex and the importance of cortical inhibition. Neuron, 38(5):689–699, 2003. ISSN 0896-6273. doi: 10.1016/s0896-6273(03)00332-5.

Yuelin Shi, Dasheng Bi, Janis K. Hesse, Frank F. Lanfranchi, Shi Chen, and Doris Y. Tsao. Rapid concerted switching of the neural code in the inferotemporal cortex. Nature, pages 1–10, 2026. ISSN 0028-0836. doi: 10.1038/s41586-026-10267-3.

Eero P Simoncelli and Bruno A Olshausen. NATURAL IMAGE STATISTICS AND NEURAL REPRESENTATION. Annual Review of Neuroscience, 24(1):1193–1216, 2001. ISSN 0147-006x. doi: 10.1146/annurev.neuro.24.1.1193.

Karen Simonyan and Andrew Zisserman. Very deep convolutional networks for large-scale image recognition. arXiv, 2014. doi: 10.48550/arxiv.1409.1556.

Yasuko Sugase, Shigeru Yamane, Shoogo Ueno, and Kenji Kawano. Global and fine information coded by single neurons in the temporal visual cortex. Nature, 400(6747):869–873, 1999. ISSN 0028-0836. doi: 10.1038/23703.

Keiji Tanaka. Inferotemporal cortex and object vision. Annual Review of Neuroscience, 19(1):109–139, 1996. ISSN 0147-006x. doi: 10.1146/annurev.ne.19.030196.000545.

Imran Thobani, Javier Sagastuy-Brena, Aran Nayebi, Jacob Prince, Rosa Cao, and Daniel Yamins. Model-brain comparison using inter-animal transforms. arXiv, 2025. doi: 10.48550/arxiv.2510.02523.

Leslie G. Ungerleider and Mortimer Mishkin. Two cortical visual systems. In [“Ingle, \ Goodale, David J. and \,\ Mansfield Melvyn A. and, and Richard J. W.”], editors, Analysis of Visual Behavior, pages 549–586. MIT Press, Cambridge, MA, 1982.

Jasper JF van den Bosch, Tal Golan, Benjamin Peters, JohnMark Taylor, Mahdiyar Shahbazi, Baihan Lin, Ian Charest, Jö rn Diedrichsen, Nikolaus Kriegeskorte, Marieke Mur, and Heiko H Schütt. A python toolbox for representational similarity analysis. 2025. doi: 10.7554/elife.107828.1.

Will Xiao, Kasper Vinken, and Margaret Livingstone. Response dynamics in macaque ventral stream recapitulate the visual hierarchy. bioRxiv, page 2025.11.11.686115, 2025. doi: 10.1101/2025.11.11.686115.

Daniel L K Yamins and James J DiCarlo. Using goal-driven deep learning models to understand sensory cortex. Nature Neuroscience, 19(3):356–365, 2016. ISSN 1097-6256. doi: 10.1038/nn.4244.

Daniel L. K. Yamins, Ha Hong, Charles F. Cadieu, Ethan A. Solomon, Darren Seibert, and James J. DiCarlo. Performance-optimized hierarchical models predict neural responses in higher visual cortex. Proceedings of the National Academy of Sciences, 111(23):8619–8624, 2014. ISSN 0027-8424. doi: 10.1073/pnas.1403112111.

Jeffrey M. Yau, Anitha Pasupathy, Scott L. Brincat, and Charles E. Connor. Curvature processing dynamics in macaque area v4. Cerebral Cortex, 23(1):198–209, 2013. ISSN 1047-3211. doi: 10.1093/cercor/bhs004.

Chengxu Zhuang, Siming Yan, Aran Nayebi, Martin Schrimpf, Michael C. Frank, James J. DiCarlo, and Daniel L. K. Yamins. Unsupervised neural network models of the ventral visual stream. Proceedings of the National Academy of Sciences, 118(3):e2014196118, 2021. ISSN 0027-8424. doi: 10.1073/pnas.2014196118.

